# Disagreement among Genomic Markers Profoundly Influences Phylogenetic Inference in Squamates

**DOI:** 10.64898/2026.01.06.698024

**Authors:** Marc Tollis, John H. Neddermeyer, Simone M. Gable

## Abstract

Phylogenomic-scale studies of the same clades using different markers and methods often support highly confident yet different species trees, obscuring many phylogenetic relationships and hampering comparative studies. Among reptiles, the >11,000 extant species of squamates (Order Squamata: lizards, snakes, and amphisbaenians) comprise a highly diverse and well-studied clade, yet many unresolved questions about squamate origins remain, including the root of the squamate phylogeny and the relationships of snakes to other toxicoferans. To understand biological, molecular, and methodological sources of phylogenomic heterogeneity in squamates, we analyzed four genome-scale marker datasets, including thousands of protein-coding genes, anchored hybrid enrichment loci, and ultra-conserved elements. We applied standardized alignment, filtering, and tree-building methods across marker sets, analyzed patterns in substitutional saturation and codon positions, and measured gene tree-species tree discordance across all data partitions. We found that many contentious relationships in squamate phylogenetics are driven by conflicts stemming from input data quality, incorrect model fit, and sampling bias, and that biological drivers of heterogeneity include both incomplete lineage sorting and introgression. We account for these sources of heterogeneity and strengthen resolution of the squamate phylogeny. Using a simulation approach, we find that current ultra-conserved element datasets for squamates deviate the most from phylogenetic expectations among marker types. Our work demonstrates that identifying sources of phylogenomic heterogeneity while accounting for input data limitations can resolve phylogenetic conflicts.

## Introduction

Phylogenomic analysis aims to use genome-scale data to understand evolutionary relationships and resolve key branches on the tree of life (Dunn et al. 2008; Kawahara et al. 2019). Despite progress towards this goal for many taxa, phylogenomic inference can still yield unresolved tree topologies, even when hundreds or thousands of markers are used (Mirarab et al. 2024; Stiller et al. 2024; Wu et al. 2024). These conflicts, or phylogenomic discordance, often take the form of (1) differences between genealogical patterns of individual loci and the true branching order of species (i.e., “gene tree–species tree discordance” (Kumar et al. 2012; Smith et al. 2015), or (2) disparate results in tree topologies from separate phylogenomic studies of the same taxa based on different marker types and tree methods (Jarvis et al. 2014; Reddy et al. 2017). Understanding the varied sources of phylogenomic conflict, and how these sources interact with the properties of different marker classes, would allow molecular systematists to account for biases and make informed decisions about which markers are most likely to resolve nodes of interest.

Biological sources of phylogenomic discordance include incomplete lineage sorting (ILS), which is when ancestral polymorphisms cause gene trees to differ from species trees (Avise and Robinson 2008; Degnan and Rosenberg 2009), and introgression, which arises when hybridization or gene flow between species drives discordance (Hibbins and Hahn 2021). These processes may be further shaped by variation in genome architecture such as recombination rate heterogeneity, which can influence genealogical signal (Mirarab et al. 2024) and allow the introgression of non-sister taxa to persist (Li et al. 2019; Burbrink et al. 2025). Recurrent substitutions at the same sites over the course of molecular evolution can cause genetic saturation, which erodes phylogenetic signal over time, and makes older divergences difficult to resolve (Philippe et al. 1994; Philippe et al. 2011). Other sources of phylogenomic discordance stemming from molecular evolutionary processes are compositional heterogeneity and codon usage bias (Cooper et al. 2003; Foster 2004; Cox et al. 2014).

Many methodological factors influence phylogenomic conflict as well, including taxonomic sampling, marker choice, alignment errors, data trimming and filtering, and model misspecification (Cooper et al. 2003; Foster 2004; Cox et al. 2014). For example, loci that are shorter or gap-rich may provide insufficient information for accurate gene tree estimation, while uneven taxon coverage can amplify noise or generate misleading topologies. Although these impacts of discordance have been documented in several clades (Singhal et al. 2021; Gable et al. 2022; Stiller et al. 2024; Morales-Briones et al.), few studies have explicitly compared how the biological, molecular, and methodological factors of discordance differ across marker types, nor the degree to which discordance across marker types affects gene tree and species tree estimation of the same clade.

Our goal in this study is to investigate sources of phylogenomic discordance across different marker types in squamates and to determine marker features that affect phylogenomic conflict for the group. Squamates are a diverse and globally-distributed reptile clade comprising >11,000 extant species of lizards and snakes, including seven major clades (Gekkota, Dibamia or Family Dibamidae, Scincoidea, Laterata/Lacertoidea, Iguania, Anguimorpha, and Serpentes) that mostly diverged during the Jurassic Period (reviewed in Simões et al. 2025). The evolutionary history of squamates involves several rapid radiations and extensive phenotypic convergence and has been studied using fossils, comparative methods, and genomics (Simões et al. 2018; Simões, Tollis, et al. 2025). Many key aspects in squamate evolutionary history remain unresolved, including (1) the position of the root of the squamate phylogeny, (2) the relationships among toxicoferan clades (Iguania, Anguimorpha, and Serpentes), and (3) several radiations within Iguania, Anguimorpha, Elapoidea, Henophidia, and Colubroidea. We summarize historically contentious nodes in the squamate phylogeny in Table 1.

**Table 1.** Major historically contentious regions of the squamate phylogeny identified across phylogenomics studies, the conflicting hypotheses proposed, representative references, and brief explanations of key issues.

| Phylogenetic Region | Conflicting Hypotheses | Representative Studies | Notes/Key Issues Identified across Studies |
| --- | --- | --- | --- |
| Root of Squamata | 1. Gekkota as sister to all other squamates<br>2. Dibamidae as sister to all other squamates<br>3. Gekkota + Dibamidae as the earliest diverging lineage | Vidal & Hedges 2005; Pyron et al. 2013; Reeder et al. 2015; Burbrink et al. 2020; Singhal et al. 2021; Title et al. 2024 | Saturation in short loci; taxon sampling differences; uneven representation of dibamids; high gene tree error in UCEs; morphological vs. molecular conflict |
| Toxicofera relationships (Iguania, Anguimorpha, Serpentes) | 1. Iguania + Serpentes<br>2. Iguania + Anguimorpha<br>3. Anguimorpha + Serpentes | Fry et al. 2006; Pyron et al. 2013; Reeder et al. 2015; Burbrink et al. 2020; Singhal et al. 2021; Mount & Brown 2022; Gable et al. 2023 | High ILS; possible ancient introgression; rapid radiation with short internal branches; marker-class sensitivity; missing data effects; conflicting gene tree distributions |
| Pleurodonta (iguanian lizards) | Relationships among major families (e.g., Liolaemidae, Iguanidae, Tropiduridae) unresolved; alternative placements of Liolaemidae | Townsend et al. 2011; Pyron et al. 2013; Esquerré et al. 2022; Singhal et al. 2021 | Reticulation in some lineages (e.g., Liolaemidae); patchy taxonomic sampling; heterogeneity in alignment quality |
| Elapoidea (advanced snakes) | Alternative placements of Pseudoaspidae, Elapidae, Lamprophiidae | Figueroa et al. 2016; Burbrink et al. 2020; Das et al. 2024 | Short, saturated loci in UCEs; low signal-to-noise ratios; unresolved backbone relationships in some analyses |
| Henophidia (boas + pythons + allies) | Unstable relationships among major henophidian families; conflict between molecular datasets and morphology | Pyron et al. 2013; Burbrink et al. 2020; Singhal et al. 2021 | Deep divergences with chronic saturation; limited genomic representation for some lineages; potential introgression signals |
| Colubroides (most extant snakes) | Conflicts among backbone relationships, especially within Colubroidea and Lamprophiidae | Figueroa et al. 2016; Burbrink et al. 2020; Singhal et al. 2021 | High gene tree error; heterogeneous alignment quality; incomplete representation of rapidly diversifying lineages |
| Placement of Xenosauridae (Anguimorph lizards) | Unstable placement across studies (sometimes sister to Anguidae, sometimes nested differently within Anguimorpha) | Reeder et al. 2015; Burbrink et al. 2020; Singhal et al. 2021 | Sensitivity to alignment and filtering choices; conflicting signals among marker types; limited taxonomic coverage |
| Placement of Shinisauria (Anguimorph lizards) | Unstable placement across studies (sometimes sister to Varanoidea, sometimes sister to Neoanguimorpha) | Fry et al. 2006; Pyron et al. 2013; Reeder et al. 2015; Burbrink et al. 2020; Singhal et al. 2021 | Limited genomic representation for some lineages; potential introgression signals |

Molecular studies differ in the placement of snakes relative to iguanians and anguimorphs, and several of these studies support a Iguania + Anguimorpha clade, with the exclusion of snakes (Vidal and Hedges 2005; Fry et al. 2006; Pyron et al. 2013; Streicher and Wiens 2017; Burbrink et al. 2020; Singhal et al. 2021; Title et al. 2024). Multiple total-evidence studies combining molecular and morphological characters have also inferred snakes as sister to Iguania + Anguimorpha (Reeder et al. 2015; Pyron 2017; Simões et al. 2018; Simões, Brum, et al. 2025). Meanwhile, empirical studies examining gene tree heterogeneity in squamate phylogenomic datasets have also revealed support for Iguania + Serpentes (Burbrink et al. 2020; Singhal et al. 2021; Gable et al. 2023), and likelihood ratio comparisons across datasets favor this topology in some analyses (Mount and Brown 2022). Similar patterns of discordance occur in other regions of the squamate tree, particularly the relationships of dibamids to gekkotans and whether they comprise the outgroup to all other squamates, as well as several other squamate radiations including within colubroid and henophidian snakes, pleurodont iguanians, and anguimorphs (Figueroa et al. 2016; Streicher and Wiens 2017; Burbrink et al. 2020; Singhal et al. 2021) (Table 1).

Phylogenomic studies attempting to resolve these conflicts in squamate systematics have used a broad range of marker types, including hundreds of anchored hybrid enrichment loci (AHEs) (Lemmon et al. 2012; Burbrink et al. 2020; Singhal et al. 2021; Title et al. 2024), thousands of ultra-conserved elements (UCEs) (Faircloth et al. 2012; Streicher and Wiens 2017; Singhal et al. 2021; Title et al. 2024), and large sets of protein-coding genes (Townsend et al. 2011; Pyron et al. 2013; Singhal et al. 2021; Gable et al. 2023; Title et al. 2024). Each marker class has distinct and documented strengths and weaknesses in their abilities to recover phylogenetic signal. UCEs provide many loci but tend to be short, gap-rich, and highly variable in informativeness (Faircloth et al. 2012; Portik and Wiens 2021). Unusually high amounts of UCE-derived gene tree errors have motivated some authors to emphasize concatenated supermatrix tree-building results over coalescent-consistent species tree approaches for squamates (Title et al. 2024). Compared to UCEs, AHE datasets for squamates have offered longer and therefore more informative loci, yet with much smaller numbers of loci (Lemmon et al. 2012; Burbrink et al. 2020; Singhal et al. 2021; Title et al. 2024). Protein-coding genes can number in the thousands and provide many informative sites for squamates and other reptiles, particularly when whole genomes are used; however, until recently large scale protein-coding gene datasets for reptiles have been difficult to obtain (Gable et al. 2022; Gable et al. 2023). Due to methodological differences across all these studies, including marker type, models used, and taxonomic sampling, determining the relative strengths and weaknesses of marker types for resolving contentious splits in the squamate phylogeny has remained difficult.

Here, we apply a standardized framework to examine patterns of phylogenomic conflict across thousands of loci and millions of DNA base pairs across three major marker classes commonly used for squamates (and vertebrate molecular systematics in general): protein-coding genes, AHEs, and UCEs. By aligning, trimming, filtering, and analyzing all datasets under controlled methods, and explicitly testing for biological (ILS, introgression), molecular (substitutional saturation, codon bias), and methodological (alignment quality, taxon sampling, model fit) sources of discordance, we isolate many factors that strongly influence phylogenetic conflict. We further use simulations to compare observed discordance against expectations under fitted substitution models to evaluate marker performance relative to null phylogenetic expectations. Specifically, we investigate (1) how marker type, data quality, model fit, and tree-building method shape discordance across the phylogeny; (2) the relative roles of ILS and introgression at unresolved nodes; (3) the extent to which accounting for saturation and alignment quality clarifies the placement of key groups such as dibamids and toxicoferans; and (4) whether some marker types are more reliable than others for resolving long-standing conflicts in squamate systematics. This comparative, standardized approach provides a generalizable framework for evaluating sources of phylogenomic discordance in other challenging clades.

## Materials and Methods

### Data sources and taxon sampling

We assembled a large protein-coding genes dataset for squamates to be used as a benchmark to compare to previously published marker datasets. We downloaded 111 squamate genome assemblies analyzed in Gable et al. (2024), representing 35 squamate families, with scaffold N50 >10kbp (except for *Dibamus bouretti*, see below), plus five outgroup taxa: tuatara (*Sphenodon punctatus*), painted turtle (*Chrysemys picta*), American alligator (*Alligator mississippiensis*), chicken (*Gallus gallus*), and human. We collected the nucleotide and amino acid sequences for single copy orthologs extracted from each assembly using BUSCO v5.4.4 (Manni et al. 2021) with the OrthoDB v10 sauropsida database (Kriventseva et al. 2019). We used sequence IDs to extract and organize orthologs into 7,480 multi-species FASTA files for evolutionary analysis following Gable et al. (2022). For the other marker sets, we downloaded published data including 370 AHEs from Burbrink et al. (2020) (Burbrink_AHEs hereafter), 388 AHEs from Title et al. (2024) (Title_AHE hereafter), and 5,023 UCEs from Title et al. (2024).

Summaries of all pre- and post-trimming datasets in terms of number of markers, species, and total base pairs (bp) are in Table 2.

**Table 2.**
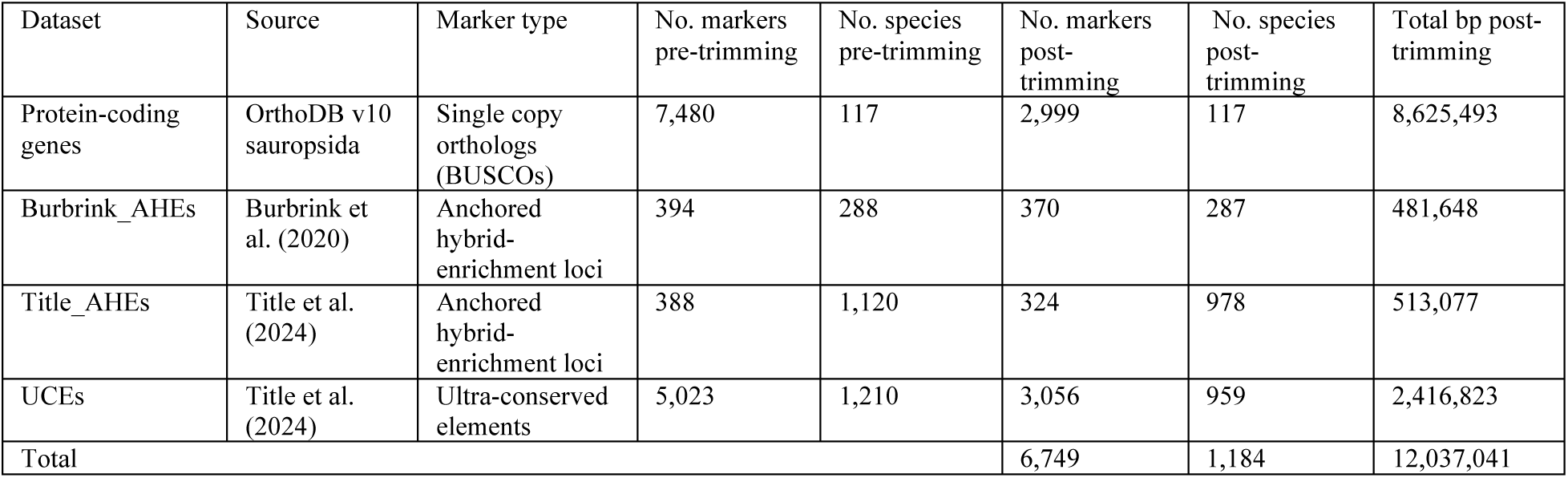
Summaries of four squamate phylogenomics datasets analyzed in this study.

| Dataset | Source | Marker type | No. markers pre-trimming | No. species pre-trimming | No. markers post-trimming | No. species post-trimming | Total bp post-trimming |
| --- | --- | --- | --- | --- | --- | --- | --- |
| Protein-coding genes | OrthoDB v10 sauropsida | Single copy orthologs (BUSCOs) | 7,480 | 117 | 2,999 | 117 | 8,625,493 |
| Burbrink_AHEs | Burbrink et al. (2020) | Anchored hybrid-enrichment loci | 394 | 288 | 370 | 287 | 481,648 |
| Title_AHEs | Title et al. (2024) | Anchored hybrid-enrichment loci | 388 | 1,120 | 324 | 978 | 513,077 |
| UCEs | Title et al. (2024) | Ultra-conserved elements | 5,023 | 1,210 | 3,056 | 959 | 2,416,823 |
| Total |  |  |  |  | 6,749 | 1,184 | 12,037,041 |

### Alignment and filtering

We aligned 7,480 protein-coding gene FASTA files with MAFFT v7.505 (Katoh and Standley 2013), using TrimAL v1.4.rev15 (Capella-Gutierrez et al. 2009) to remove all positions in alignments with gaps in 30% or more of sequences (-gt 0.7). Summary statistics for all alignments including number of taxa, alignment length, missing data percentage, number and proportion of variable and parsimony-informative sites, and GC content, were estimated with AMAS v3.04 (Borowiec 2016). We removed trimmed protein-coding gene alignments <1500 bp in length and containing >15% gaps and >80% taxa representation to maximize informativeness of protein-coding genes in reptiles (Karin et al. 2020; Gable et al. 2022).

To standardize alignment quality across UCE and AHE datasets, we first trimmed all downloaded alignments with TrimAL using options -resoverlap 0.7 -seqoverlap 70 -noallgaps to set minimum overlap and percentage, and to remove columns composed of only gaps. We then realigned the trimmed data using MAFFT v7.505 with option “—auto”, followed by re-trimming using a 30% gap threshold (-gt 0.7). We pruned unidentifiable sequences and dropped species for which there were <50 markers represented per dataset, which is the minimum recommended for confident phylogenetic inference using summary-based coalescent species tree methods (Ruane et al. 2015) (Supplementary Table 1). We realigned the pruned alignments with MAFFT v7.505 (option “—auto”).

### Phylogenetic inference

We inferred maximum likelihood gene trees for each dataset with IQ-TREE v2.2.0.4 (Minh, Schmidt, et al. 2020), using ModelFinder (Kalyaanamoorthy et al. 2017) and determined the best-fit model for each partition. For the protein-coding genes and AHEs marker sets, we used the resulting gene trees to infer a coalescent-consistent multilocus species tree with ASTRAL-III (Zhang et al. 2018), with branch support measured in local posterior probabilities.

We used a slightly different ASTRAL tree method for the UCE gene trees to avoid unrealistic computational runtimes. We measured branch support for all UCE gene trees with the SH-like approximate likelihood ratio test (aLRT) in IQ-TREE, followed by species tree inference with WASTRAL v1.18.3.5 (Zhang and Mirarab 2022), which uses a hybrid weighting scheme based on aLRT and branch length. After initial UCE-based tree visualizations of gene and species trees, we found evidence of contamination and unknown/improbable sequence IDs and pruned 14 additional taxa from UCE alignments and gene trees. We regenerated the UCE-based species tree from the pruned data using WASTRAL.

We concatenated the protein-coding genes, AHE, and UCE alignments into separate partitioned supermatrices and inferred maximum likelihood species trees for each marker set with IQ-TREE MPI multicore v2.2.2.7. We performed model testing on the protein-coding genes and UCE supermatrices in IQ-TREE with ModelFinder and PartitionFinder2 (Lanfear et al. 2017) with options “-m TESTMERGEONLY -rclusterf 20” to consider the top 20% partition schemes using the fast relaxed clustering algorithm for improved computational efficiency, and merge partitions if warranted. We then used options “--redo-tree -st NT -bb 1000 -alrt 1000” to resume tree reconstruction with the specified models and partitions with 1,000 ultrafast bootstrap replicates and the Shimodaira-Hasegawa-like (SH-like) approximate likelihood ratio test (Shimodaira 2002). AHE supermatrix trees were run with option “-m TESTMERGE” to conduct model testing in ModelFinder and PartitionFinder2 to merge partitions if warranted, along with 1,000 ultrafast bootstrap replicates and the SH-like approximate likelihood ratio test.

### Heterogeneity analyses

We collapsed all tree branches to family-level using a custom Python script with packages ete3 (Huerta-Cepas et al. 2016) and pandas (The pandas development team 2024), following taxonomic classification reported in the Reptile Database (Uetz et al. 2025). For squamate families with only one taxon in a tree, we renamed the tip with the family name.

For each marker set, we calculated pairwise Robinson-Foulds (RF) distances (Robinson and Foulds 1981) between gene trees and their species tree(s) using multiRF in phytools (Revell 2024), and pairwise normalized RF distances using the RF.dist from phangorn (Schliep 2011). We additionally calculated RF distances among species trees inferred from different marker classes to quantify cross-marker topological discordance. We computed gene and site concordance factors for all branches on the full ASTRAL and concatenated supermatrix trees for each dataset in IQ-TREE (Minh, Hahn, et al. 2020).

The gene concordance factor (gCF) is the proportion of gene trees whose topologies are consistent with every node in the species tree, and gDF1 and gDF2 are the proportions of the second and third most common gene tree topologies (Minh, Hahn, et al. 2020). To test for evidence of introgression events relative to ILS at each split in the squamate species trees, we compared the frequencies of gDF1 and gDF2 under the null expectation that under ILS gDF1 and gDF2 should be roughly equal, and introgression is detectable when one minor gene tree topology is more frequent (Green et al. 2010; Minh, Hahn, et al. 2020). We used a chi-square test to evaluate the null hypothesis at each node, adjusting for multiple testing using a false discovery rate 0.05. Nodes rejecting the ILS-only null were interpreted as having support for introgression.

### Substitutional saturation and codon usage bias analyses

We estimated levels of substitutional saturation for all four marker-type datasets. We first calculated the slopes of the linear regressions between the raw pairwise and TN93-corrected distances for a subset of 822 protein-coding genes with ≥95% species coverage, 324 Title_AHEs, 3,056 Title_UCEs, and 370 Burbrink_AHEs using the R package ape (Paradis et al. 2004). Unsaturated alignments should have a slope of 1, while saturated alignments should have a slope approaching zero (Philippe et al. 1994). We categorized alignments above the mean slope as “unsaturated” and below the mean as “saturated”. To investigate the effects of substitutional saturation on species tree estimation, we inferred species trees in ASTRAL-III using gene trees from saturated and unsaturated alignments. We used the multi-threaded WASTRAL by Branch Length (mode 3) for UCEs. We computed the saturation ratio for each taxonomic family as (saturated/(saturated + unsaturated)).

To further understand saturation effects, we re-aligned 822 protein-coding gene alignments with ≥95% species sampling (to focus on high-confidence codons) using codon-aware MACSE v11.05 (Ranwez et al. 2011), concatenated and partitioned by gene and codon position with AMAS, and inferred maximum likelihood phylogenies for saturated and unsaturated (1) first and second and (2) third codon positions in IQ-TREE with model testing and 1,000 ultrafast bootstrap replicates. We also constructed a maximum likelihood tree using a supermatrix of MACSE amino acid alignments with model testing in IQ-TREE and inferred an ASTRAL tree from amino acid gene trees.

To test for evidence of codon usage bias in our protein-coding genes dataset, we estimated the relative synonymous codon usage bias (RSCU) of the 822 MACSE alignments using CodonW v1.4.4 (Peden 1999). The RSCU measures how frequent a codon is observed relative to its expected frequency in the absence of any codon usage bias. The RSCU indices were then used in a factor analysis in the FactoMineR v2.11 R package (Lê et al. 2008).

### Evaluation of overall marker performance relative to simulated data

One definition of marker performance is the ability of a marker to capture evolutionary relationships despite inherent and common biological and methodological drivers of phylogenomic discordance. Marker performance in this sense has been evaluated by comparing empirical data to simulations under reasonable baseline phylogenetic expectations (Borowiec et al. 2025), although few studies have compared marker performance across different markers from the same clade. We compared observed and simulated phylogenomic discordance across marker types to determine which marker type performs closest to null expectations.

First, we concatenated and partitioned supermatrices and estimated a maximum likelihood guide tree for each marker type in IQ-TREE, applying a separate GTR+F+I+gamma substitution model to all partitions. We then calculated observed gene concordance factors for each marker type using gene trees estimated with the same substitution model in IQ-TREE. To generate simulated marker sets representing null phylogenetic expectations, we used substitution model parameters estimated from each observed alignment. We simulated alignments using AliSim (Ly-Trong et al. 2022) in IQ-TREE (iqtree2 –alisim -s -t) with the observed maximum likelihood tree as the guide for each marker set. For each simulation, we generated a custom guide tree by pruning the observed species tree to match taxon sampling of the observed alignment. We generated gene trees from the simulated alignments in IQ-TREE under the same substitution model (iqtree2 -s -alrt 1000 -m) and calculated simulated gene concordance factors on the observed species tree.

We compared observed and simulated gene concordance factors using linear regression. Because the protein-coding genes dataset contained fewer nodes, we tested for a small-N effect by comparing the distribution of slopes and intercepts from 1,000 AHE and UCE subsets randomly down sampled to 100 nodes.

## Results

### Alignment quality, informativeness, and species tree topologies differ sharply among marker types

We analyzed a large number of sequences, gene trees, and species trees in our study were, and while all datasets received an equal amount of scrutiny, many intermediate results were a recapitulation of previous studies (Burbrink et al. 2020; Singhal et al. 2021; Gable et al. 2024; Title et al. 2024), as expected. Therefore, we first present the protein-coding gene results, which are new to our study. Additional figures for all datasets are in the Supplementary Material, and their results are discussed below. All datasets and scripts for analyzing the results will be made publicly available on Dryad upon acceptance. After trimming and filtering for length and species representation, we obtained 2,999 protein-coding gene markers from 111 publicly available squamate assemblies representing 35 squamate families and all seven major clades, plus five outgroup taxa (Table 1). On average, 86% of known single-copy BUSCOs were present in the assemblies. Alignments averaged 2,786 bp in length with ∼92% taxonomic coverage, ∼1,745 (60%) parsimony-informative sites, and 3.7% gaps (Fig. 1a-d, Supplementary Fig. 1).

**Figure 1.**
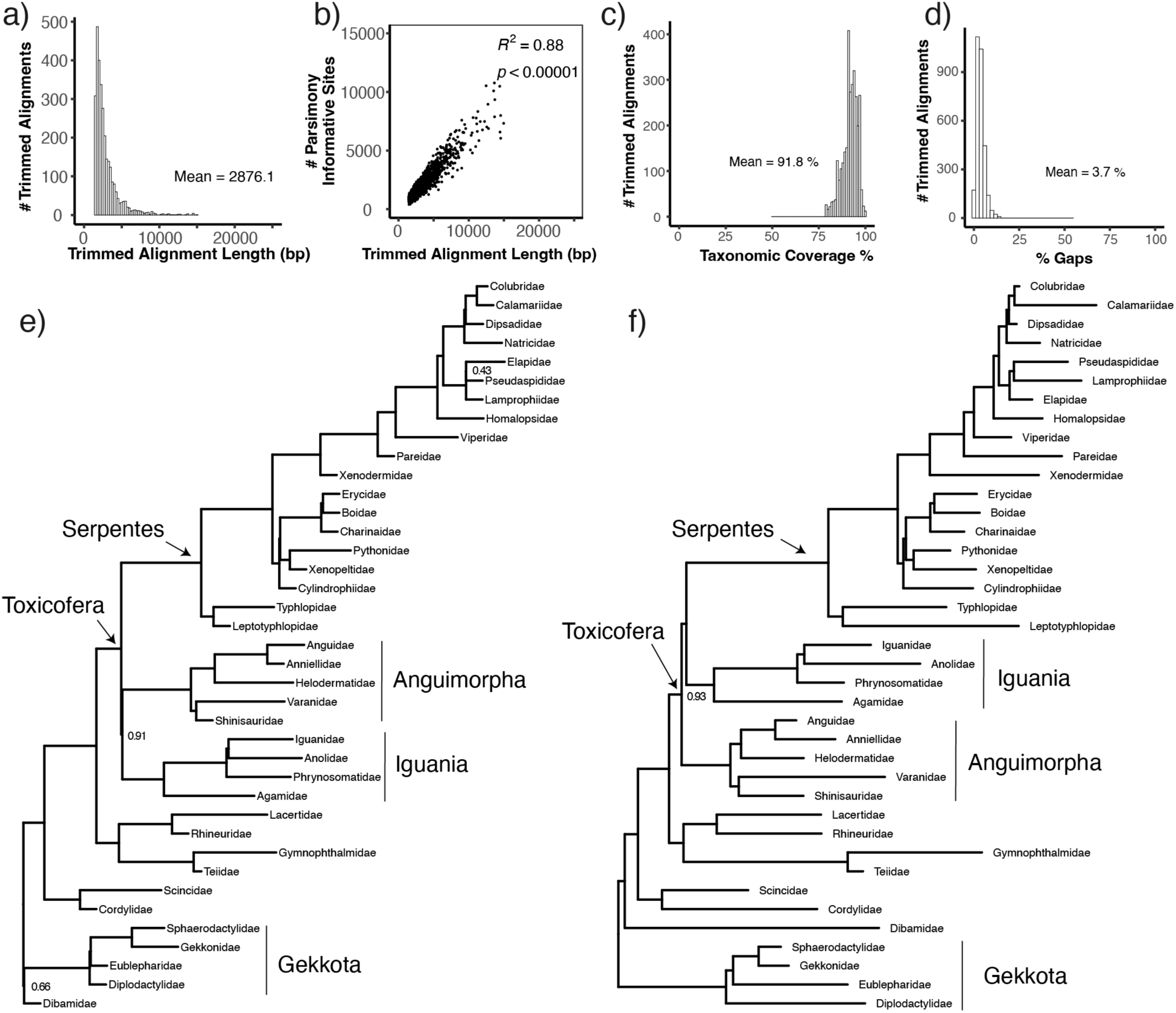
Data quality metrics and inferred species trees from 2,999 protein-coding gene (BUSCO) alignments for squamates. Length distribution (a), correlation between length and informativeness (b), distribution of taxonomic coverage (c), and distribution of percent gaps (d) for 2,999 trimmed protein-coding gene alignments. Family-level species trees (e,f) marking the placement of toxicoferans and gekkotans with respect to dibamids, inferred using the gene trees in ASTRAL (e), and with maximum likelihood using a concatenated supermatrix (f). All branches received 100% support (local posterior probability and bootstrap value, respectively) except where indicated.

The protein-coding gene ASTRAL tree was strongly supported overall (97% branches with posterior probability or pp > 0.95), and support was lower at historically problematic nodes (Table 1, Fig. 1e, Supplementary Fig. 2). A branch shared by Dibamidae and Gekkota formed the outgroup to all other squamates (pp =0.66). Within Toxicofera, Iguania and Anguimorpha were sister taxa with the exclusion of Serpentes (pp = 0.91). Additionally, family-level positions within the snake superfamily Elapoidea (Pseudaspididae and Elapidae) had lower support (pp = 0.43). The protein-coding genes supermatrix tree placed gekkotans as the outgroup to all other squamates, followed by the divergence of dibamids (bootstrap support or bs =100%), and supported an Iguania + Serpentes clade (bs = 100%) (Fig. 2f; Supplementary Fig. 2).

**Figure 2.**
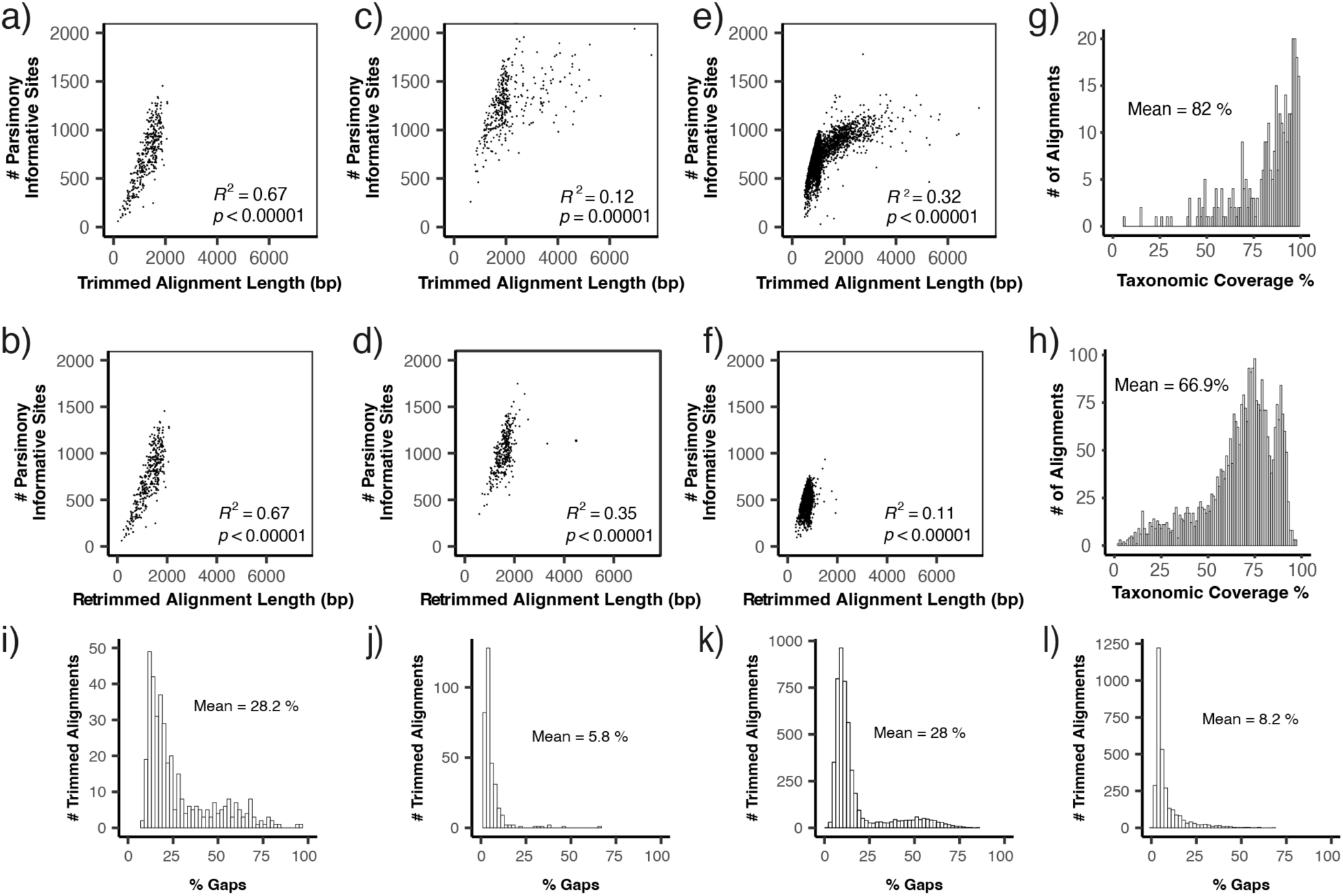
Data analysis of three sets of squamate phylogenomic alignments. Trimmed and retrimmed Burbrink_AHEs were highly informative (a,b). Retrimming improved informativeness for Title_AHEs (c,d), but not for UCEs (e,f). Overall taxonomic coverage for Title_AHEs (g) was higher than for UCEs (h). Retrimming improved gaps for Title_AHEs (i,j) and UCEs (k,l).

After trimming and pruning, we obtained 370 Burbrink_AHEs, 324 Title_AHEs, and 3,056 UCEs (Table 1). Trimming and retrimming analyses revealed distinct data-quality profiles for the two AHE datasets as well as for the UCEs (Fig. 2, Supplementary Fig. 3). Retrimmed Burbrink_AHEs retained high taxonomic coverage (92% per locus) and strong informativeness (Fig. 2a,b). Informativeness was improved markedly after retrimming the Title_AHEs (Fig. 2c,d), but not UCEs (Fig. 2e,f). While considerably more abundant (3,056 loci), UCEs had much lower taxonomic coverage (Fig. 2g,h) and were more gap-rich, although retrimming improved gaps overall (Fig. 2i-l). Final trimmed Title_AHE and UCE alignments lacked representation for Dibamidae due to low species coverage (<50 markers).

### Widespread topological conflict in squamate phylogenetic reconstruction

All ASTRAL and concatenated supermatrix species trees for Burbrink_AHEs, Title_AHEs, and UCEs are provided in the Supplementary Material (Supplementary Fig. 4, Supplementary Fig. 5, Supplementary Fig. 6). Like the protein-coding genes ASTRAL tree, the Burbrink_AHEs ASTRAL tree weakly supported a Dibamidae + Gekkota clade at the root of the squamate phylogeny (pp= 0.65). However, the Burbrink_AHEs supermatrix tree supported Dibamidae as sister to all non-gekkotan squamates (bs = 79%). There were several conflicts regarding the placement of the toxicoferan clades (Iguania, Anguimorpha, and Serpentes) across AHE and UCE datasets and methods as well. Besides the protein-coding genes supermatrix tree, all species trees recovered an Iguania + Anguimorpha clade within Toxicofera with varying levels of bootstrap support across supermatrices (30.5%-100%) and 100% posterior probabilities across ASTRAL results. Overall, RF distances reflected greatest dissimilarity between UCE-derived species trees and all other marker sets, with more agreement between protein-coding genes and both AHE datasets.

There were many other areas of conflict in our results across datasets and tree methods (Fig. 2e-f, Supplementary Fig. 4, Supplementary Fig. 5, Supplementary Fig. 6). These include differences in the estimated relationships of pleurodont iguanians, with low branch support throughout, and the phylogenetic positions of several snake families within Colubroides also conflicted between tree methods and datasets. Other known areas of conflict across methods included Booidea, Elapoidea, Colubroidea, the placement of Eublepharidae within Gekkota, and the placement of Xenosauridae in Anguimorpha.

Robinson-Fould’s distance analysis also revealed stark differences in gene tree topologies with respect to species tree topology for each data type (Supplementary Fig. 7; Supplementary Fig. 8). Mean normalized Robinson-Fould’s distance between gene trees and species trees was lowest for protein-coding genes (0.25), followed by Burbrink_AHEs (0.32), Title_AHEs (0.35), and highest for UCEs (0.47). Robinson-Fould’s distance analysis also revealed extensive topological conflicts in squamate phylogeny across datasets and across ASTRAL and supermatrix species tree reconstruction methods (Fig. 3a-d). Overall, protein-coding genes showed the least disagreement between the two tree methods (Fig. 3a).

**Figure 3.**
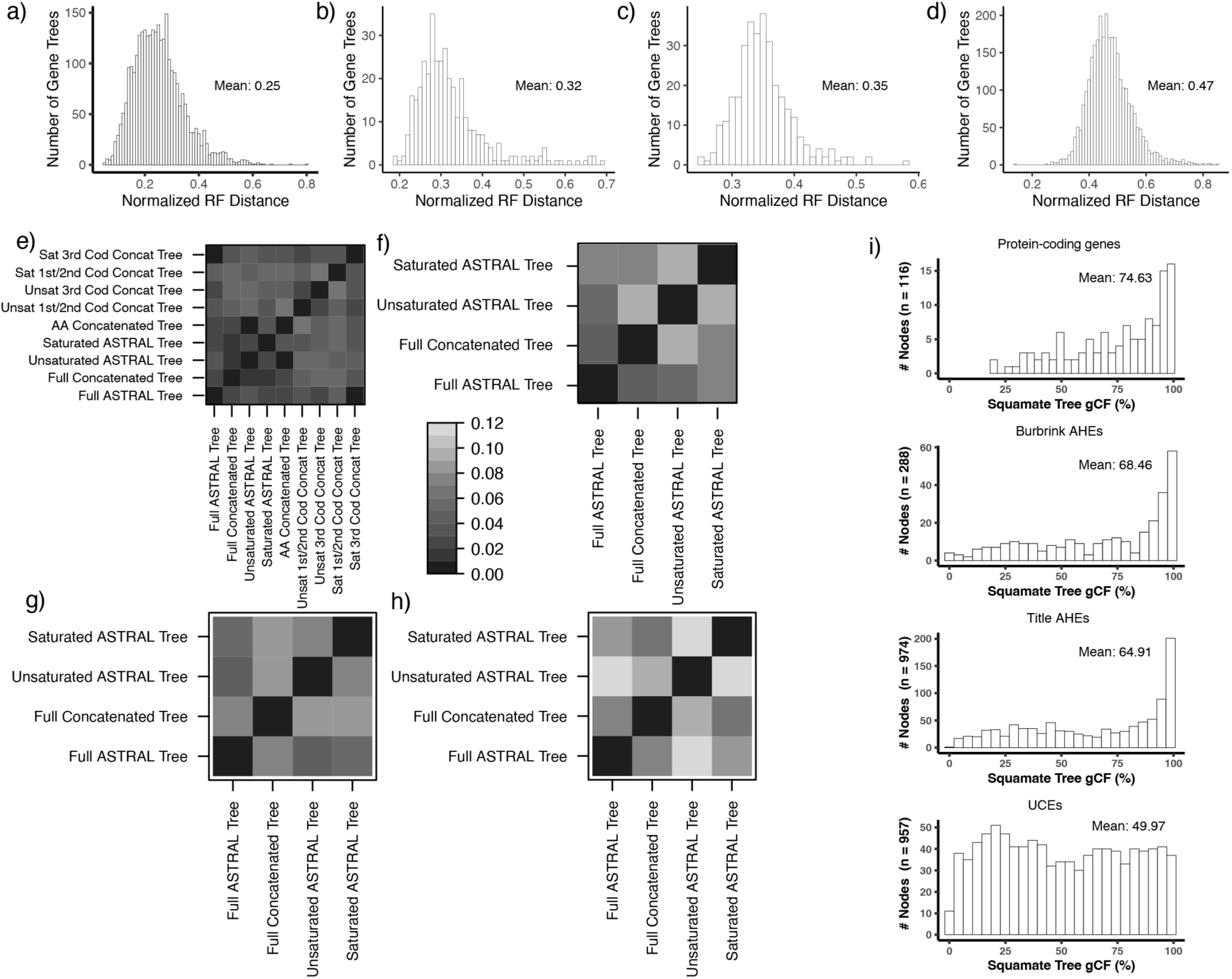
Topological conflict in squamate phylogenomics across tree methods and markers. Normalized Robinson-Fould’s (RF) distances calculated between gene trees and species trees for protein-coding genes (a), Burbrink_AHEs (b), Title_AHEs (c), and UCEs (d). RF distances between species-level species trees (ASTRAL, concatenation) revealed the most consistent agreement from protein-coding genes (e), followed by Burbrink_AHEs (f), Title_AHEs (g) and lastly UCEs (h). RF distances for family-level trees are in Supplementary Fig. 8. Gene concordance factor (gCF%) analyses comparing topologies of gene trees and species trees revealed the highest concordance from protein-coding genes and the lowest concordance from UCEs (i). Site concordance factors are in Supplemental Fig. 9.

### Incomplete lineage sorting and introgression drive phylogenomic conflict in squamates

Gene and site concordance factor (gCF and sCF) analyses allowed us to compare gene tree-species tree discordance and heterogeneity across the four marker types (Fig. 3e). Overall, protein-coding genes yielded higher gCFs (mean=75%), followed by Burbrink_AHEs (mean=69%), Title_AHEs mean=65%), and lastly UCEs (mean=50%). Meanwhile, sCFs across datasets were largely uniform (mean∼54-55%) (Supplementary Fig. 9).

Our chi-square tests for evidence of introgression by comparing the frequencies of minor gene topologies yielded significant results for 33 branches for protein-coding genes, 42 branches for Burbrink_AHEs, 142 branches for Title_AHEs, and 259 branches for Title_UCEs. We did not find evidence for introgression at the root of Squamata (Table 3). In contrast, we rejected the null hypothesis at the Iguania-Anguimorph split for protein-coding genes, Title_UCEs, and Burbrink AHEs; two of these remained significant after correction. We also found support for introgression at the Shinisauridae + Varanoidea branch within anguimorphs, with the most common alternate topologies placing *Shinisaurus* with neoanguimorphs to the exclusion of Varanoidea (Table 3). We also found varying support for introgression at multiple internal nodes within Pleurodonta, Scincidae, and among henophidian snake families.

**Table 3:** Chi-square test for evidence of introgression at nodes in the squamate ASTRAL species trees describing the relationships at the root of the phylogeny, among toxicoferans, and within anguimorphs. gN = number of possible gene trees; gCF = gene concordance factor; gDF1 = proportion of gene trees supporting the first minor tree topology; gDF2 = proportion of gene trees supporting the second minor tree topology; n.s. = not significant

| Split in species tree | Dataset | gN | gCF | gDF1 | gDF2 | Most common alternate topology | Raw p-value | FDR-corrected p-value |
| --- | --- | --- | --- | --- | --- | --- | --- | --- |
| Dibamidae + Gekkota | Protein-coding genes | 302 | 32% | 26% | 28% | Gekkota at the root | n.s. | n.s. |
|  | Burbrink AHEs | 219 | 31% | 29% | 27% | Gekkota at the root | n.s. | n.s. |
| Iguania + Anguimorpha | Protein-coding genes | 2998 | 32% | 21% | 31% | Iguania + Serpentes | **<0.00001** | **<0.00001** |
|  | Burbrink AHEs | 370 | 30% | 29% | 20% | Iguania + Serpentes | **0.017** | n.s. |
|  | Title AHEs | 295 | 34% | 3% | 1% | Anguimorpha + Serpentes | n.s. | n.s. |
|  | UCEs | 2871 | 25% | 15% | 18% | Anguimorpha + Serpentes | **<0.01** | **<0.05** |
| Anguimorpha: Paleoanguimorpha (Shinisauria + Varanoidea) | Protein-coding genes | 2852 | 50% | 6% | 38% | (Shinisauria + Neoanguimorpha) + Varanoidea | **<0.00001** | **<0.00001** |

Table 3: Chi-square test for evidence of introgression at nodes in the squamate ASTRAL species trees describing the relationships at the root of the phylogeny, among toxicoferans, and within anguimorphs.
|  |  |  |  |  |  |  |  |  |
| --- | --- | --- | --- | --- | --- | --- | --- | --- |
| as sister to<br>Neoanguimorpha) | Burbrink_AHEs | 351 | 57% | 26% | 6% | (Shinisauria +<br>Neoanguimorpha) +<br>Varanoidea | **<0.00001** | **<0.00001** |
|  | Title_AHEs | 313 | 95% | 0.6% | 0.6% | (Shinisauria +<br>Neoanguimorpha) +<br>Varanoidea | n.s. | n.s. |
|  | UCEs | 335 | 29% | 10% | 20% | Shinisauria + (Varanoidea +<br>Neoanguimorpha) | **<0.001** | **<0.01** |

### Substitutional saturation and the root of the squamate phylogeny

We found similar distributions and means in saturation slopes for each dataset, with average slope ranging 0.76-0.82 (Fig. 4a-d). However, RF distances revealed that levels of substitutional saturation substantially affected species tree inference (Fig. 3a-d). For protein-coding genes (Fig. 3a) and Burbrink_AHEs (Fig. 3b), ASTRAL trees inferred from saturated data conflicted the least with supermatrix trees and conflicted most with ASTRAL trees inferred from all (including both unsaturated and saturated) alignments. Overall, more discordant UCE-derived trees were most affected by saturation, and the highest RF distances in our analysis were between unsaturated and saturated ASTRAL UCE trees (Fig. 3d), reflecting the short length and high variance in informativeness of UCE loci.

**Figure 4.**
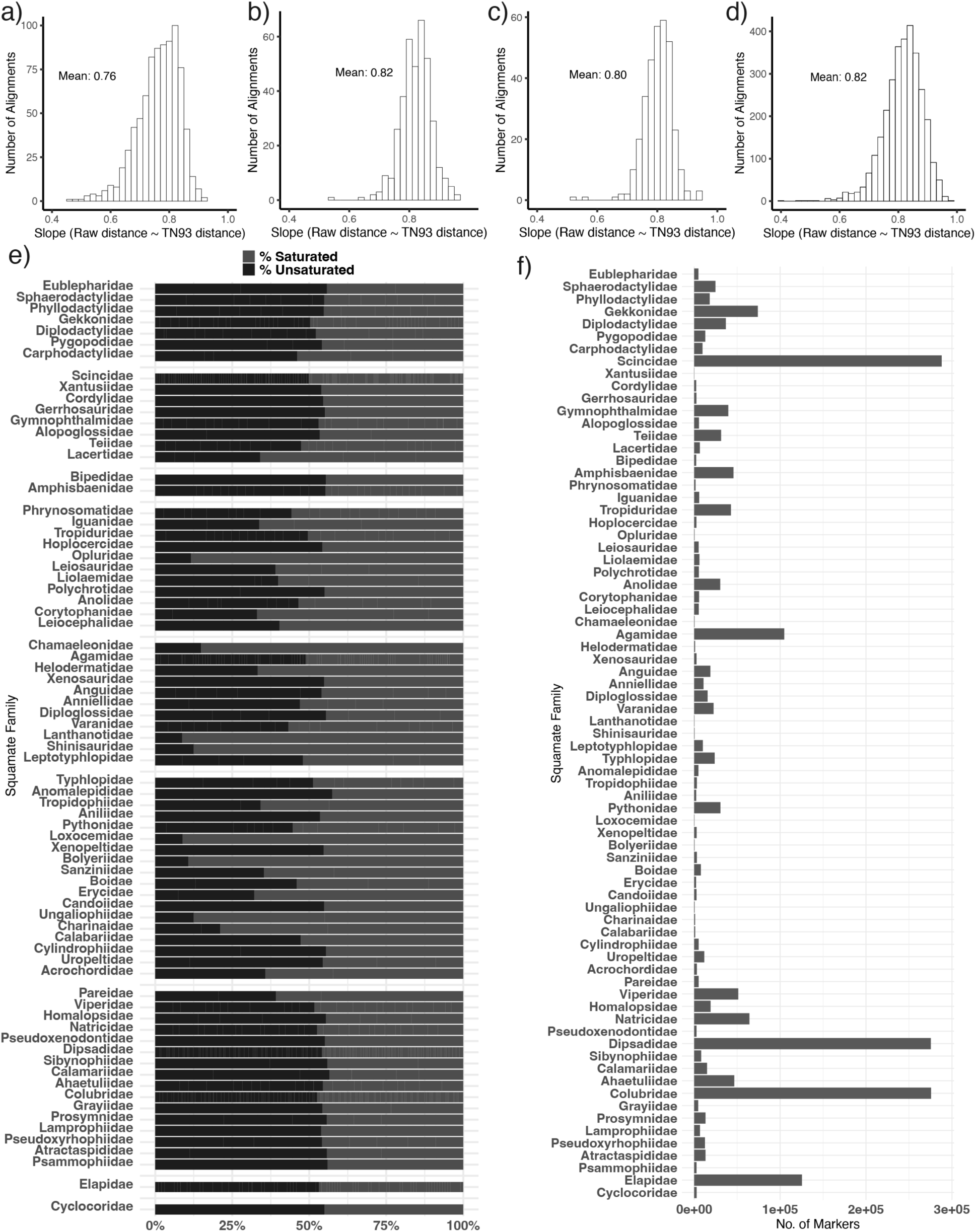
Saturation and taxonomic sampling affecting squamate phylogenetic inference. Histograms of saturation slopes across datasets: (a) BUSCO, (b) Burbrink AHEs, (c) Title AHEs, (d) UCEs. (e) Proportion of saturated vs. unsaturated markers per taxonomic family (showing lineage-level variation, especially in UCEs). (f) Total number of UCE loci for each species summed per family.

The ratios of saturated markers were similar across all datasets (51-55%) (Supplementary Fig. 10). Protein-coding genes showed the least variation in saturation ratios across squamate families, while Burbrink_AHEs showed similar consistency, although Trogonophiidae and outgroup taxon *Sphenodon* had moderately higher proportions of unsaturated alignments (∼65-70%), whereas Gerrhosauridae had a lower proportion with ∼39% of alignments unsaturated (Supplementary Fig. 11). Title_AHEs showed more variation in saturation across families, and UCEs were highly heterogenous with 67-91% saturation in some taxa, particularly pleurodont iguanians and henophidian snakes (Fig. 4e). Families with elevated saturation also tended to have lower overall UCE representation (Fig. 4f), suggesting an interaction between marker drop-out and saturation effects.

Accounting for genetic saturation strengthened the evidence for Dibamidae + Gekkota as the outgroup to all other squamates: unsaturated-only ASTRAL analyses of protein-coding genes and Burbrink_AHEs supported this arrangement with 0.89 and 0.76 bootstrap support, respectively – substantially higher than trees inferred from all loci in each of those datasets (pp=0.66 and 0.65, respectively) (Supplementary Fig. 2; Supplementary Fig. 5). Meanwhile, saturated-only ASTRAL trees supported Gekkota as the outgroup, followed by the divergence of Dibamidae (pp=0.49 for protein-coding genes, pp=0.42 Burbrink_AHEs). Further, the placement of the root of the squamate phylogeny differed across separate ASTRAL analyses of saturated and unsaturated 1^st^ and 2^nd^ as well as 3^rd^ codon positions (Supplementary Fig. 12). Together, these analyses show that inferred root placement is sensitive to saturation, consistent with expectations from deep-time phylogenetics (Lozano-Fernandez 2022).

Correspondence analyses of RSCU revealed variation in the usage of UCG, ACG, CCG, and GCG codons, but no phylogenetic bias of codon usage (Supplementary Fig. 13). The only outlier in codon usage along the second dimension was an outgroup taxon (*Homo sapiens*).

### Comparison of simulated and empirical datasets

We compared empirical versus simulated gCF values to evaluate marker behavior under baseline phylogenetic expectations. We found a strong fit between observed and simulated gene concordance factors for protein-coding genes (slope = 0.98, R^2^ = 1.0, p<0.00001) (Fig. 5). The Burbrink_AHE dataset also performed well compared to simulations (slope = 0.98, R^2^ = 0.99), as did Title_AHEs (slope = 0.98, R^2^=0.99), and Title_UCEs (slope = 1.01, R^2^ = 0.98). Residual plots supported the tightest fit between observed and fitted gCF values centered around zero for protein-coding genes, while AHEs and UCEs exhibited systematic departures from zero, and UCEs having the greatest overall residual spread (Supplementary Fig. 14). These deviations reflect higher gene-tree error and/or stronger sensitivity to model violations in AHEs and especially UCEs. The down-sampled distributions of intercepts and slopes revealed no small-N effect for protein-coding genes (Supplementary Fig. 15).

**Figure 5.**
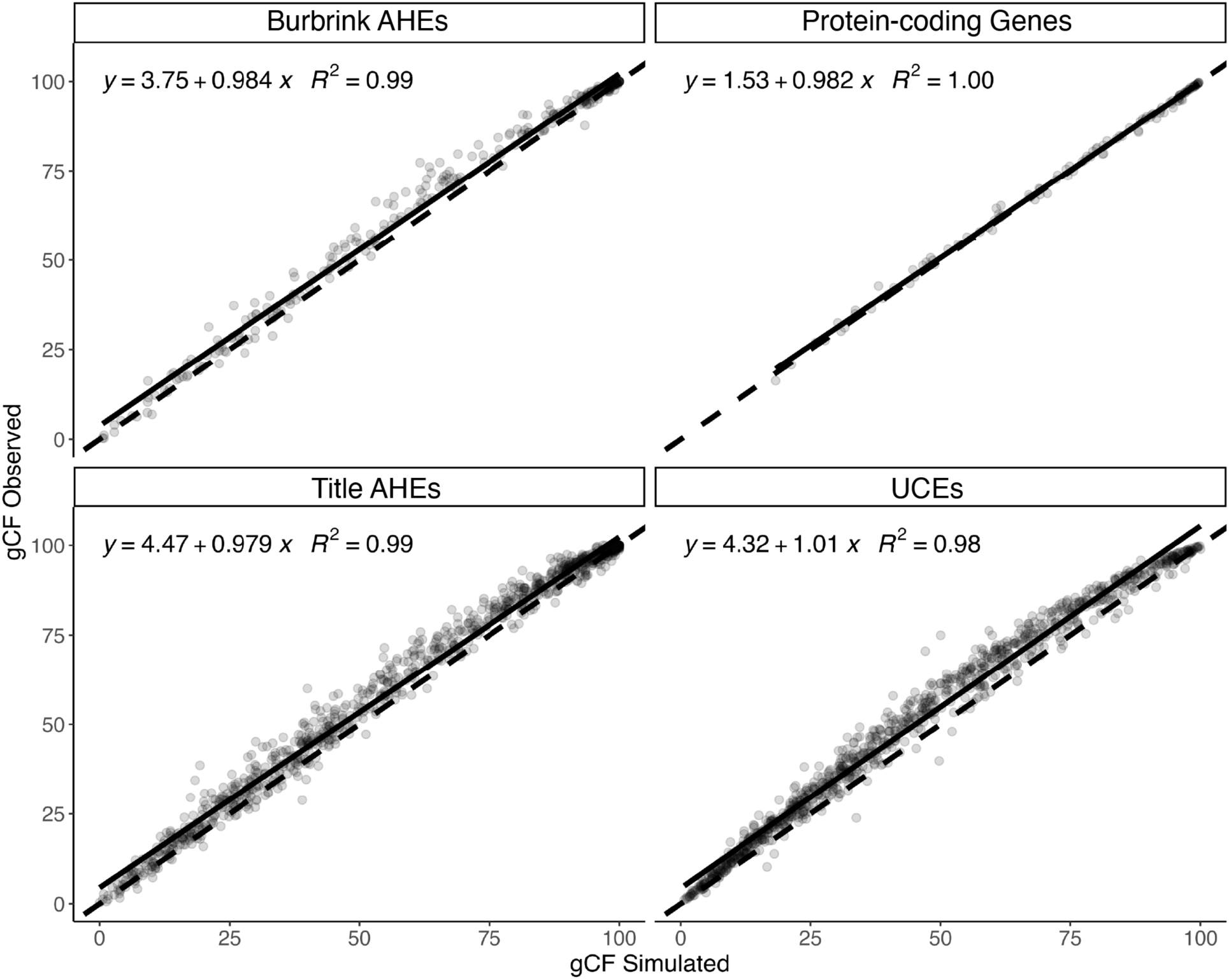
Marker performance compared to null phylogenetic expectations. Observed versus simulated gene concordance factors (gCF) for protein-coding genes, Burbrink_AHEs, Title_AHEs, and UCEs. For each marker class, simulated alignments were generated under substitution models and parameters estimated from the empirical data using the corresponding maximum-likelihood guide tree, and gCFs were calculated on the observed species tree. Solids lines indicate best-fit linear regressions, dashed lines indicate the 1:1 expectation. Slopes near unity indicate close agreement between observed and simulated gCFs under the fitted substitution model, whereas increased dispersion around the regression reflects greater deviation from model-conditional expectations. These simulations evaluate substitutional fit rather than full genealogical realism and are therefore relative benchmarks for comparing marker behavior across datasets. Residual distributions and down-sampling analyses are shown in Supplementary Figures 14 and 15.

## Discussion

### The effects of data quality, missing data, and tree methods on phylogenomic conflict in squamates

A central outcome of our study is that alignment quality and informativeness differed sharply among marker types, and these differences directly impacted phylogenetic inference in squamates. Protein-coding genes and AHEs retained high taxonomic coverage and clear phylogenetic signal after trimming and filtering, while UCEs remained short and gap-rich, with highly uneven informativeness and taxonomic representation across squamate families. Even after discarding ∼2,000 low-quality UCE alignments, the remainder still showed weak relationships between alignment length and informative sites. This is consistent with previous findings that UCEs often contain limited per-locus signal (Brown and Thomson 2017; Portik and Wiens 2021). Importantly for squamates, Title et al. (2024) reported high gene tree error and unusual species tree topologies using the same UCE dataset analyzed here, leading them to prioritize concatenated analyses. This interaction between taxonomic dropout, saturation, and short-locus signal helps explain the instability of UCE-based gene trees.

Concatenation-based and coalescent-based tree-building methods also behaved differently in our study. Concatenation inflated the bootstrap support at contentious nodes (e.g., Iguania + Serpentes), even where concordance factors were showing substantial gene tree conflict. This likely due to a sampling effect on bootstrapping that has long been recognized (Felsenstein 1985; Felsenstein and Kishino 1993; Rokas et al. 2003; Simmons and Norton 2014). For example, concatenated analysis on protein-coding genes yielded 100% bootstrap support for Iguana + Serpentes, while ASTRAL recovered Iguania + Anguimorpha with 0.91 posterior support, reflecting 32% of gene trees supporting the latter. Meanwhile, ASTRAL topologies also shifted depending on marker type and saturation partitioning, highlighting their sensitivity to gene tree error. Thus, neither concatenation nor coalescent approaches can universally resolve phylogenomic conflict in squamates, and the utility of tree-building methods depends on the signal quality and characteristics of the underlying markers. Further integration of genome-scale molecular datasets with total-evidence analyses accounting for phenotypic data type and fossil sampling may help resolve uncertainties in the deep-time relationships of squamates and other organisms (Simões, Brum, et al. 2025; Simões, Tollis, et al. 2025).

### Incomplete lineage sorting and introgression in Toxicofera and beyond

Our results suggest that while ILS has a strong influence on phylogenetic uncertainty in squamates, asymmetric minor-topology frequencies are consistent with introgression contributing at some nodes, particularly within toxicoferans. For instance, in three datasets, gDF1 and gDF2 were significantly different at the Iguania + Anguimorph species tree node, and Iguania + Snakes was the most common minor topology. This consistent asymmetry is difficult to explain with ILS alone and is compatible with deep reticulation events (Burbrink and Gehara 2018; Li et al. 2019). One explanation is that, in addition to ILS, ancient gene flow between iguanians and snakes shaped early toxicoferan divergence.

We also identified signatures of introgression in varanoids, pleurodont iguanians, scincids, and several snake lineages including henophidians. These results align with documented reticulation in Liolaemidae (Esquerré et al. 2022) and elapoid snakes (Das et al. 2024). Although some of these lineages may fall within the anomaly zone, where high levels of ILS can mimic introgression signal (Linkem et al. 2016; Singhal et al. 2021), our results indicate that a purely ILS-based explanation of their discordance is insufficient. Instead, multiple biological processes – including ILS, gene flow, and rapid divergence – interact to generate the heterogeneous signal observed across datasets. Going forward, the complexity of many squamate radiations should be more closely studied using network-aware phylogenetic approaches that can explicitly handle reticulation (Solís-Lemus and Ané 2016).

### Substitutional saturation and the root of the squamate phylogeny

Our results are consistent with genetic saturation influencing deeper nodes in the squamate phylogeny, including the placement at the root. Saturated protein-coding genes and Burbrink_AHEs recovered Gekkota at the root of squamates, whereas unsaturated partitions increased support for Dibamidae + Gekkota. This directional shift suggests that saturated loci lacking phylogenetic information are obscuring evolutionarily deeper signal in the data, paralleling other vertebrate systems where saturation reduces accuracy at ancient divergences (Philippe et al. 2011; Gable et al. 2022). Therefore, differences in the estimated roots of the squamate phylogeny across studies are likely highly to be influenced by genetic saturation, consistent with expectations for deeper phylogenetic splits.

A broader implication is that with high levels of missing data and short alignments, such as with UCEs, certain markers are more susceptible to saturation-driven errors. Our family-level analyses revealed that UCEs from poorly sampled pleurodont iguanians and henophidian snakes were disproportionally saturated compared to other markers, and these same groups showed unstable placements in UCE-based trees. Thus, uneven taxonomic sampling interacts with saturation to amplify discordance in the loci – a pattern rarely evaluated explicitly. These findings reinforce that assessing and partitioning by saturation is essential for deep time phylogenomics.

### Marker performance and future directions

To evaluate the degree to which discordance reflected phylogenetic signal versus model limitations, we compared observed and simulated gene concordance factors (gCF) for each marker set. All datasets closely matched simulation expectations; however, one key distinction across marker sets was the pattern and magnitude of the residuals from these analyses, particularly for UCEs. Protein-coding genes showed tightly centered residuals while AHEs and especially UCEs showed systematic departures. These deviations indicate that UCE discordance exceeds expectations from neutral processes alone and reflects model violations, high gene tree error, and the intrinsic limitations of short, sparsely informative loci. We note that these simulations do not incorporate ILS and therefore evaluate substitutional fit rather than full genealogical realism. Accordingly, these analyses are intended as relative, model-conditional benchmarks across marker types rather than tests of biological accuracy. Future investigations of marker performance should account for explicit sources of gene tree error, including ILS and introgression.

In addition to more simulation studies, a key future direction in phylogenomic inference for squamates will be the incorporation of genomic context – specifically the recombination landscape and physical linkage of loci. Marker performance depends partly on whether loci represent independent recombination blocks, and understanding where markers occur in the genome can help address the effects of locus non-independence on species tree estimation. For example, collapsing thousands of relatively short or uninformative UCE loci into a much smaller number of independent recombination blocks may increase informativeness and reduce gene tree error. Applying similar genomic-context filtering to squamate datasets would reduce pseudo-replication, improve coalescent inference, and clarify whether certain conflicts stem from linkage rather than biological signal. This would provide a clear path toward evaluating whether discordance arises from biological processes or from clustered, non-independent sampling of the genome.

## Conclusion

We evaluated phylogenomic discordance across four datasets representing three major marker classes – protein-coding genes, anchored hybrid enrichment loci (AHEs), and ultra-conserved elements (UCEs) – in squamates, including >8,000 loci spanning ∼12 million bp and >1,300 species. We tested how biological processes (ILS, introgression) and molecular and methodological factors (substitutional saturation, alignment quality, codon bias, model fit) contribute to conflict in squamate phylogenetics. Our results show that (1) ILS and introgression jointly shape relationships among toxicoferan clades; (2) substitutional saturation strongly influences the inferred placement of the root of the squamate phylogeny, with unsaturated loci supporting Gekkota + Dibamidae at the root; (3) UCEs are disproportionately sensitive to missing data and saturation, consistent with their high gene tree error and unusual topologies; and (4) protein-coding genes provide the most consistent phylogenetic signal for squamates under our standardized analyses and modeling assumptions, producing topologies closest to model expectations and exhibiting the lowest overall discordance. These findings help clarify the drivers of discordance in squamates and offer a general framework for evaluating marker performance in other clades where phylogenomic conflict is pervasive.

## Funding Acknowledgement

This work was supported by the National Science Foundation (grant number DEB 2323124) and a State of Arizona Technology Research Initiative Fund (TRIF) Faculty Support Grant.

## Acknowledgments

We would like to acknowledge Nicholas Bushroe and Vahid Nikoonejad Fard for assistance and helpful discussion in the early phases of this study. We also extend gratitude for Tiago Simões and Frank Burbrink for helpful commentary on drafts of the manuscript. We thank Chao Zhang for useful comments regarding tree inference. We would like to acknowledge the Monsoon computing cluster at Northern Arizona University (https://nau.edu/high-performance-computing/) for providing the computational resources necessary to carry out this study.

## Data Availability Statement

All datasets and scripts for analyzing the results will be made publicly available on Dryad upon acceptance.

## Appendices

Supplementary Materials.

