## Supplementary Figures for "Disagreement among Genomic Markers Profoundly Influences Phylogenetic Inference in Squamates"

**Supplementary Figure 1: Alignment statistics for untrimmed (left) and trimmed and filtered (right) protein-coding gene (BUSCO) alignments.**

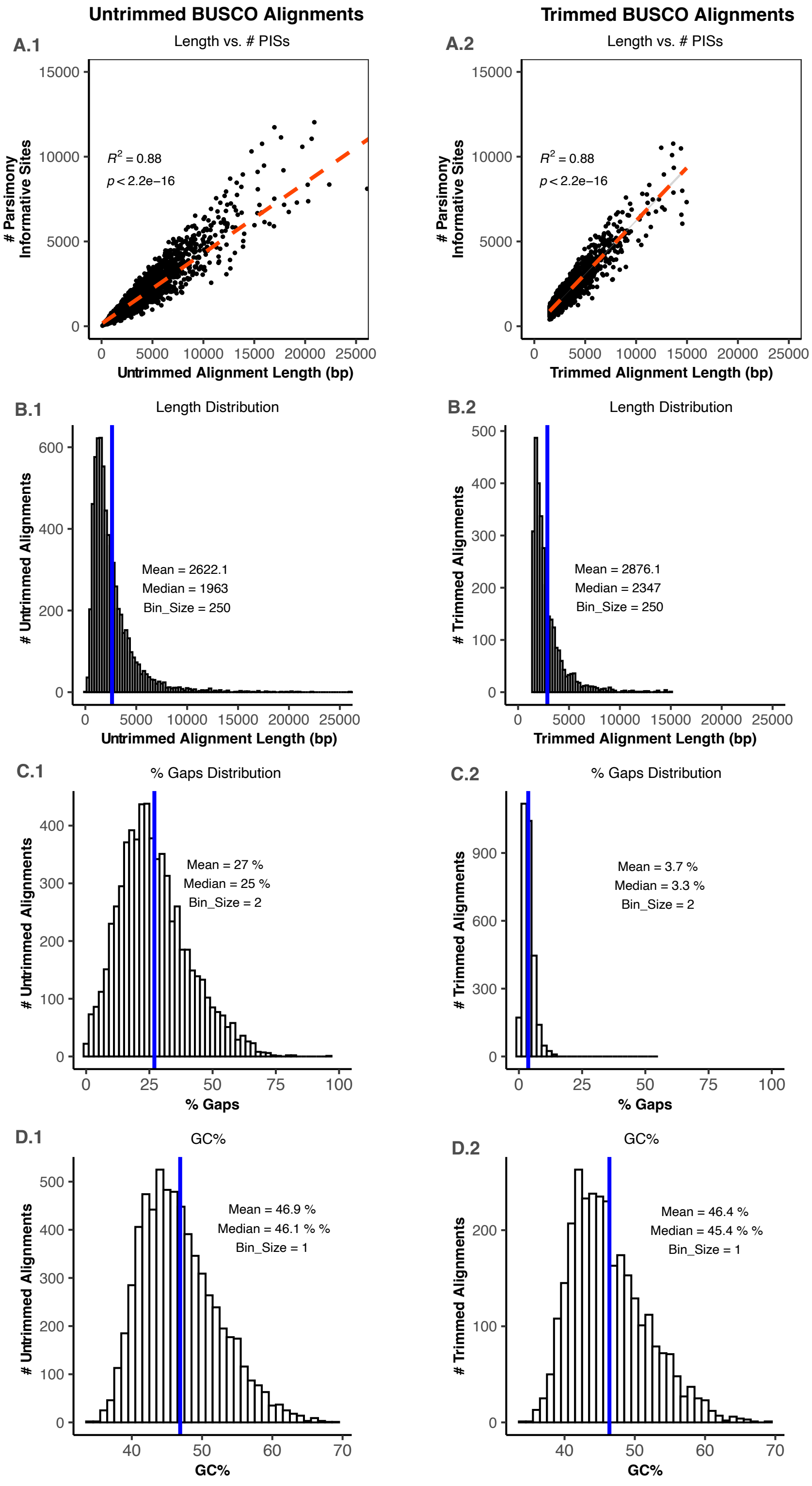

**Supplementary Figure 2: Family-level trees for protein-coding genes.** (a) ASTRAL tree, (b) unsaturated ASTRAL tree, (c) saturated ASTRAL tree, (d) concatenated supermatrix trees. ASTRAL trees use branch length scale in coalescent units. Supermatrix trees scale is in substitutions per site.

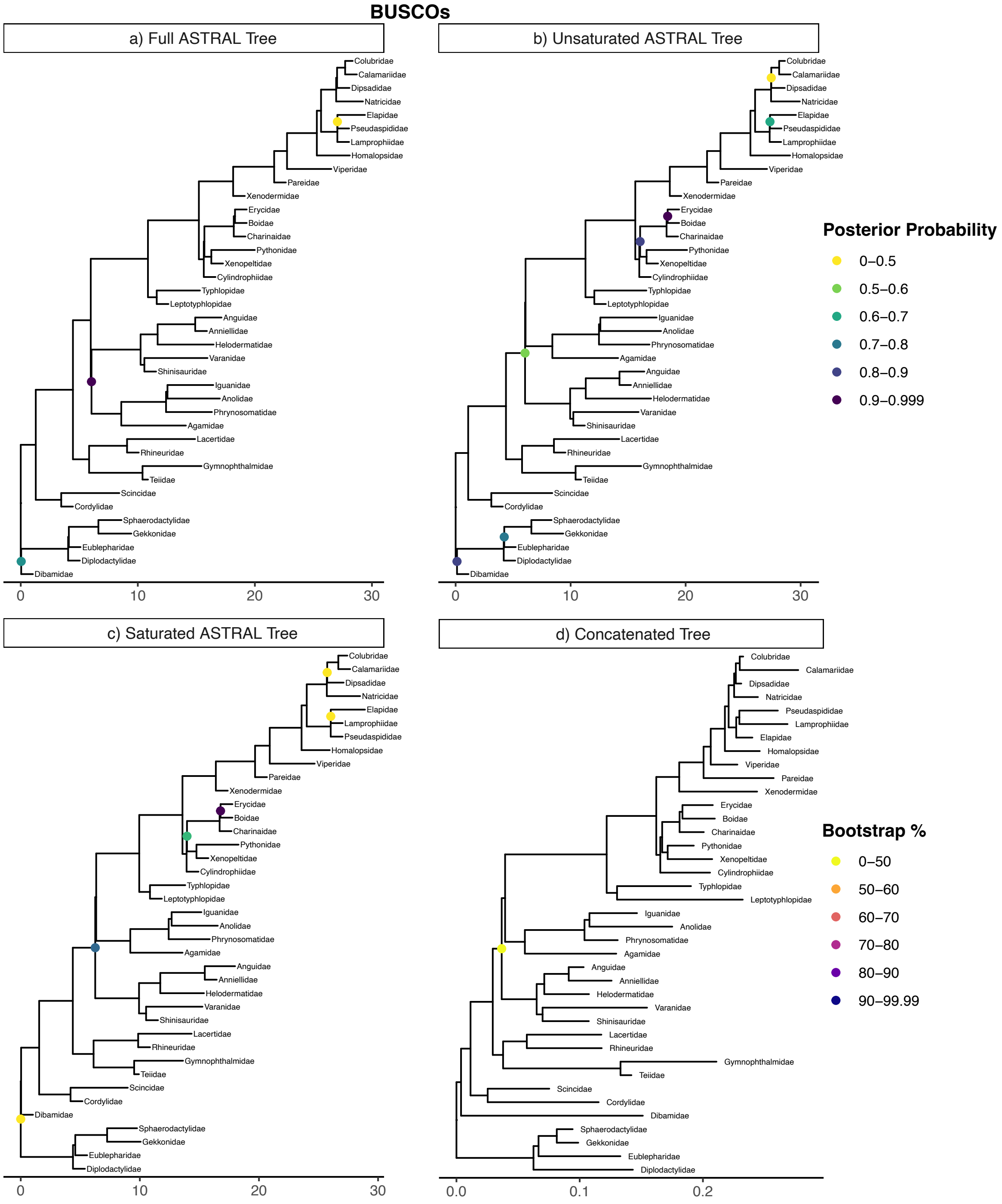

**Supplementary Figure 3: Alignment statistics across three squamate phylogenetic datasets.** Before and after refinement, labeled (A-F).

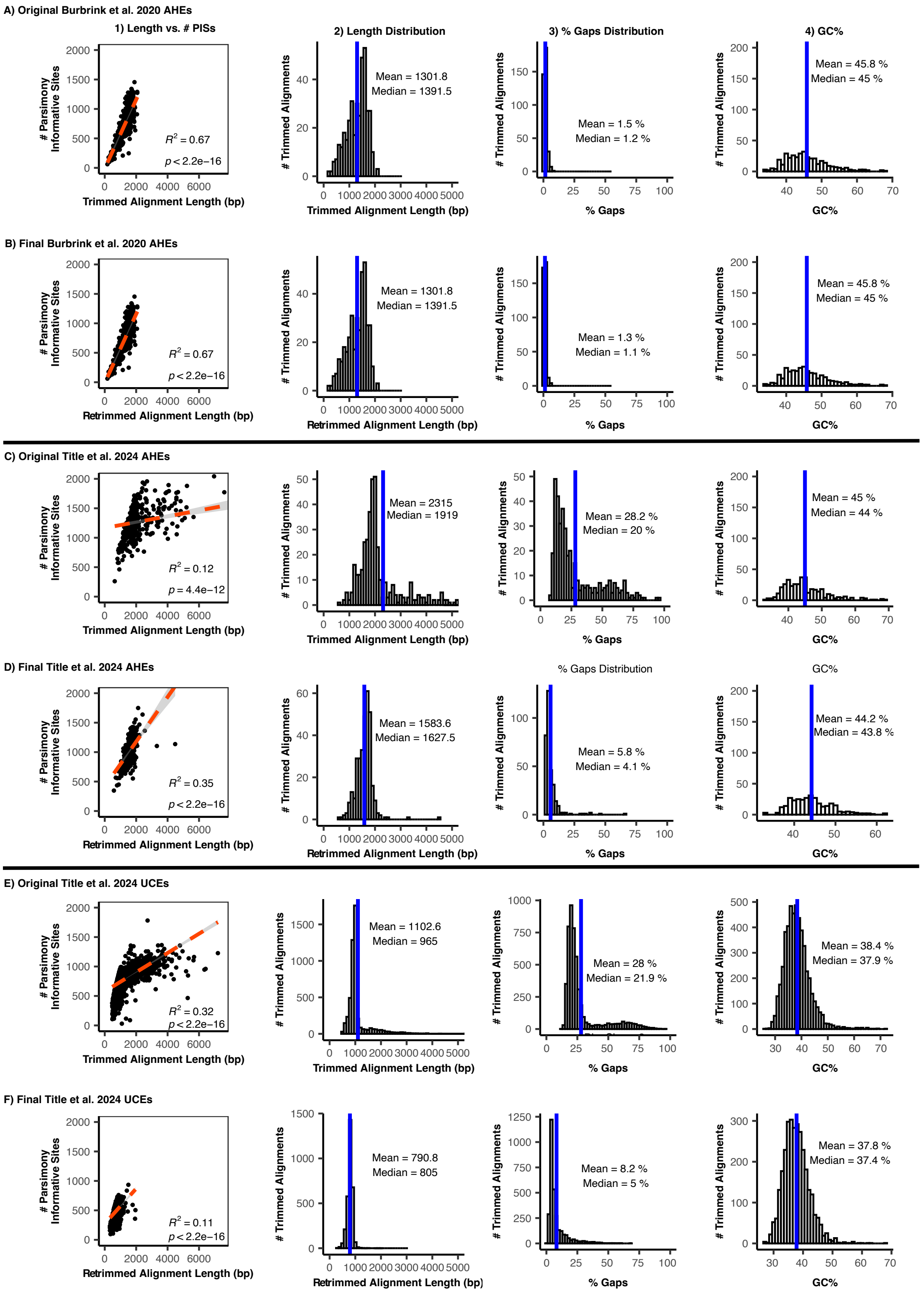

**Supplementary Figure 4: Family-level trees for Burbrink AHEs.** (a) ASTRAL tree, (b) unsaturated ASTRAL tree, (c) saturated ASTRAL tree, (d) concatenated supermatrix trees. ASTRAL trees use branch length scale in coalescent units. Supermatrix trees scale is in substitutions per site.

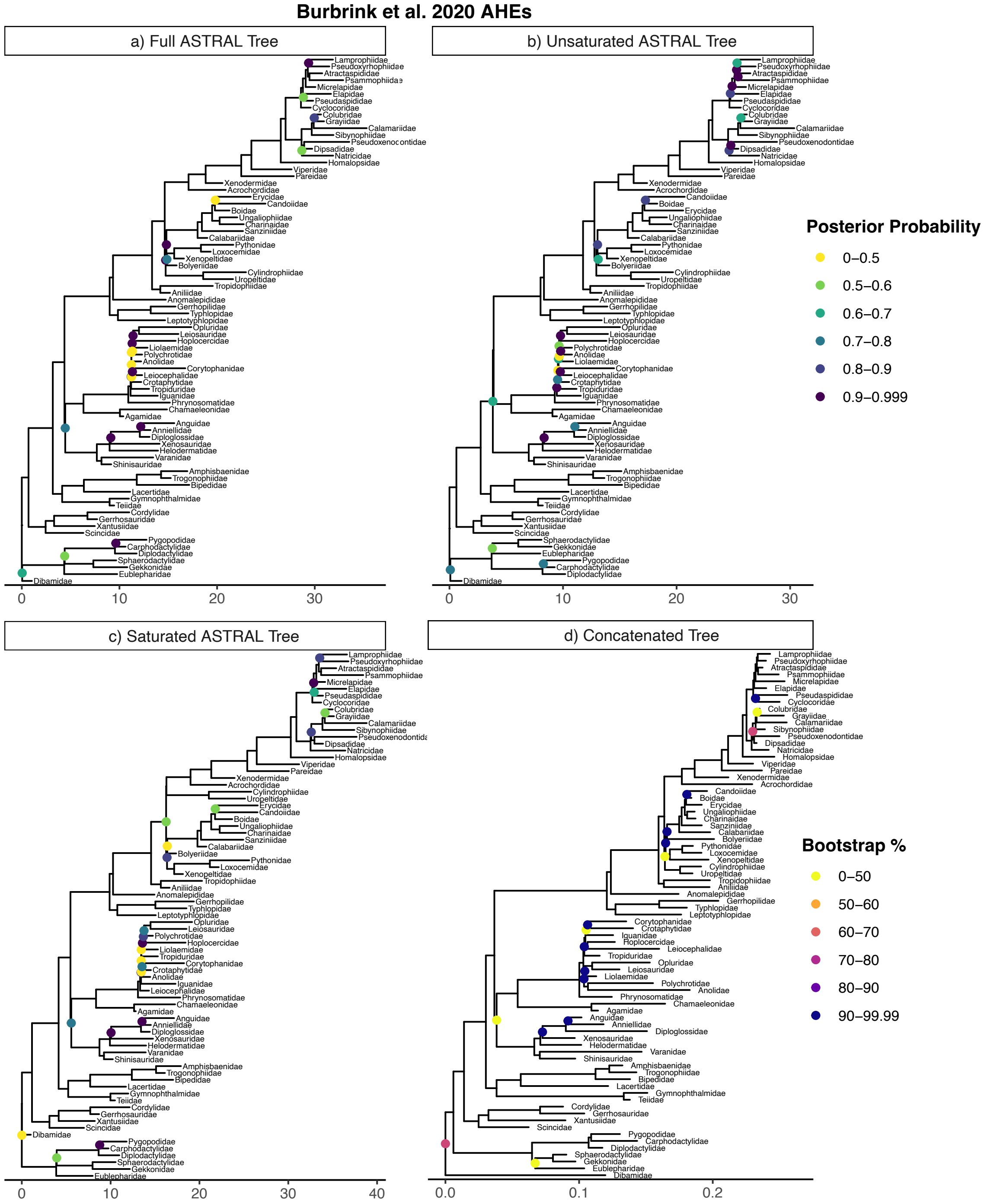

**Supplementary Figure 5: Family-level trees for Title AHEs.** (a) ASTRAL tree, (b) unsaturated ASTRAL tree, (c) saturated ASTRAL tree, (d) concatenated supermatrix trees. ASTRAL trees use branch length scale in coalescent units. Supermatrix trees scale is in substitutions per site.

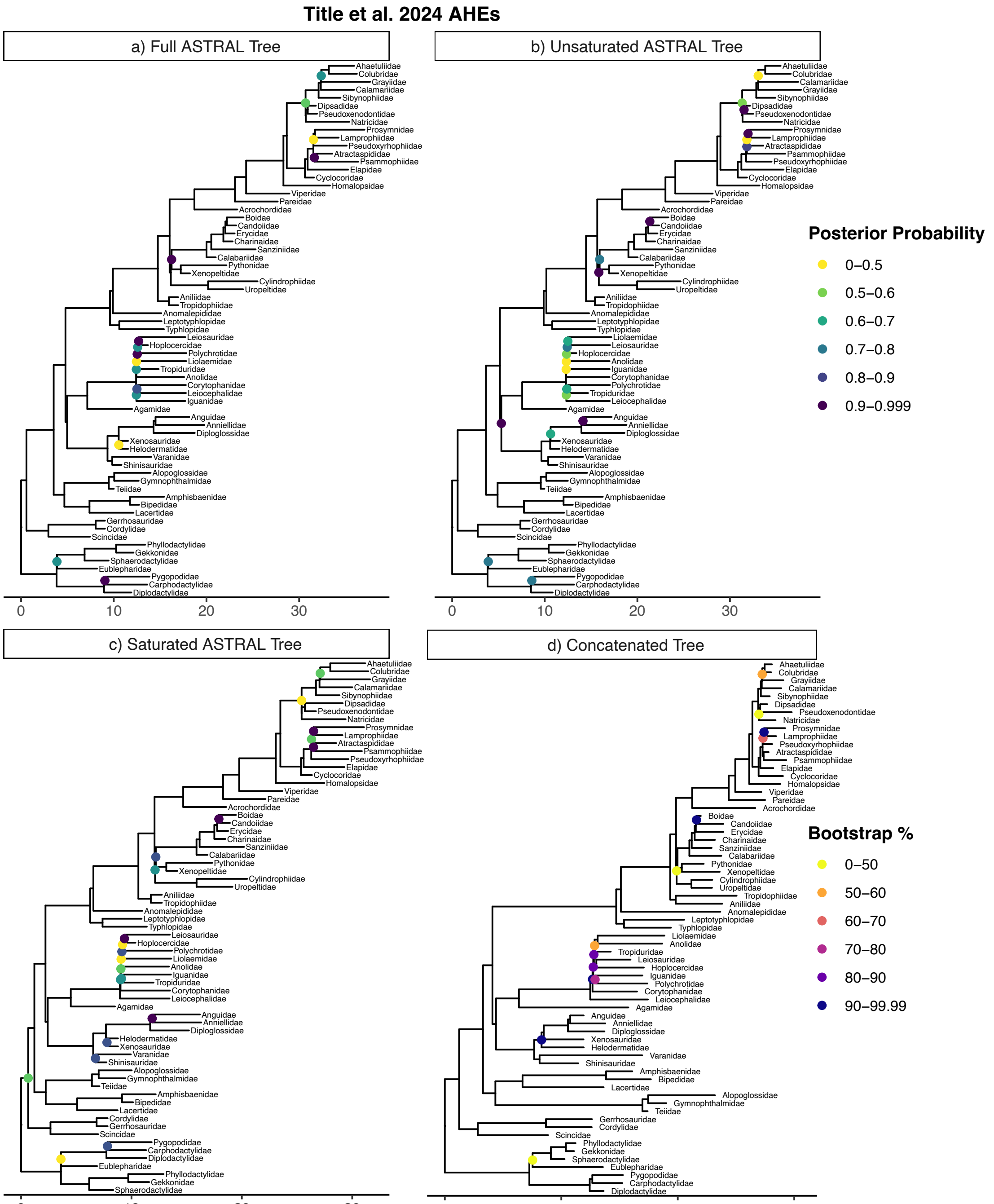

**Supplementary Figure 6: Family-level trees for Title UCEs.** (a) ASTRAL tree, (b) unsaturated ASTRAL tree, (c) saturated ASTRAL tree, (d) concatenated supermatrix tree. ASTRAL trees use branch length scale in coalescent units. Supermatrix trees scale is in substitutions per site.

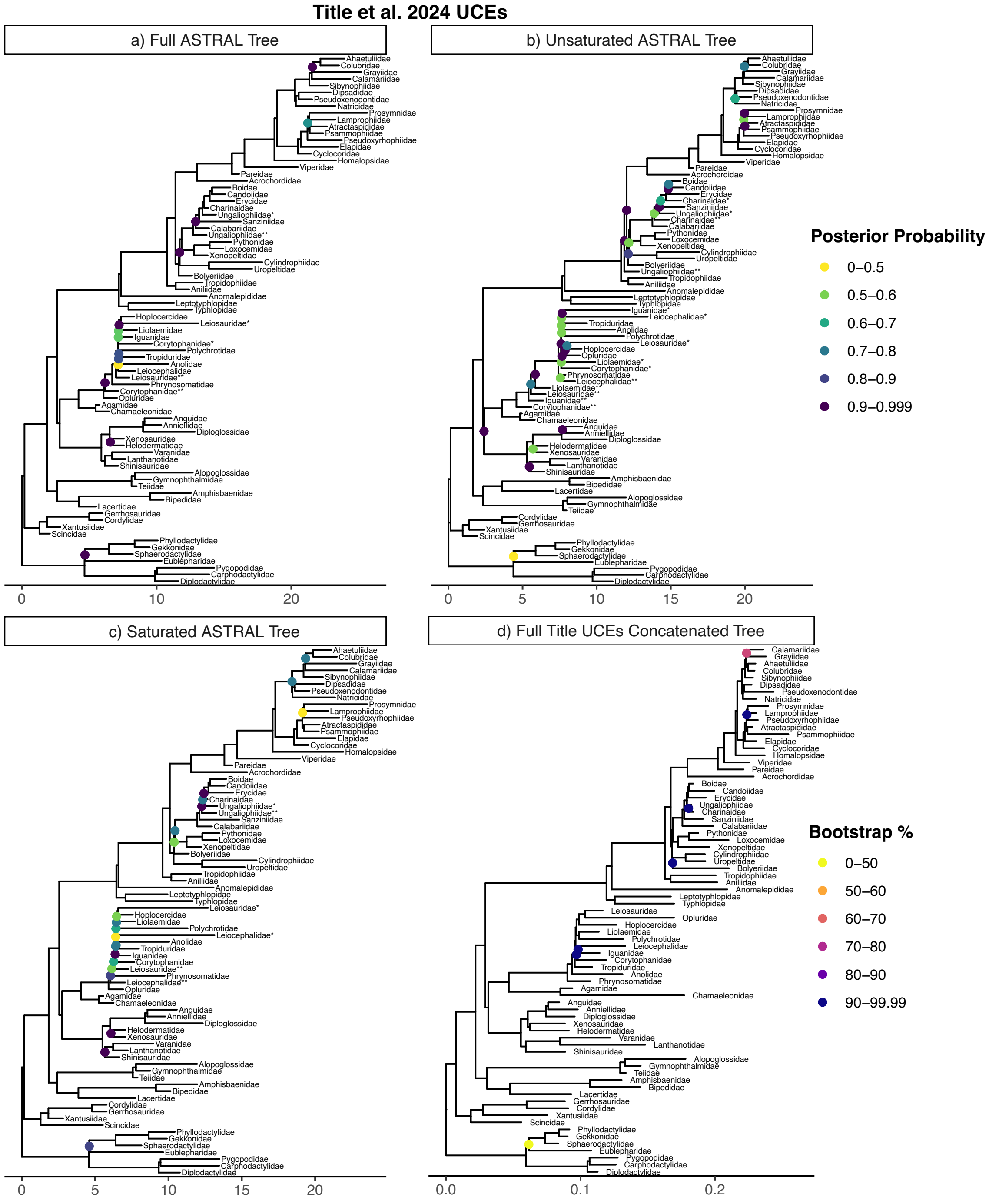

**Supplementary Figure 7: Robinson-Foulds distances across all species trees.** Comparison of topological differences between all ASTRAL- and supermatrix-inferred species trees by dataset.

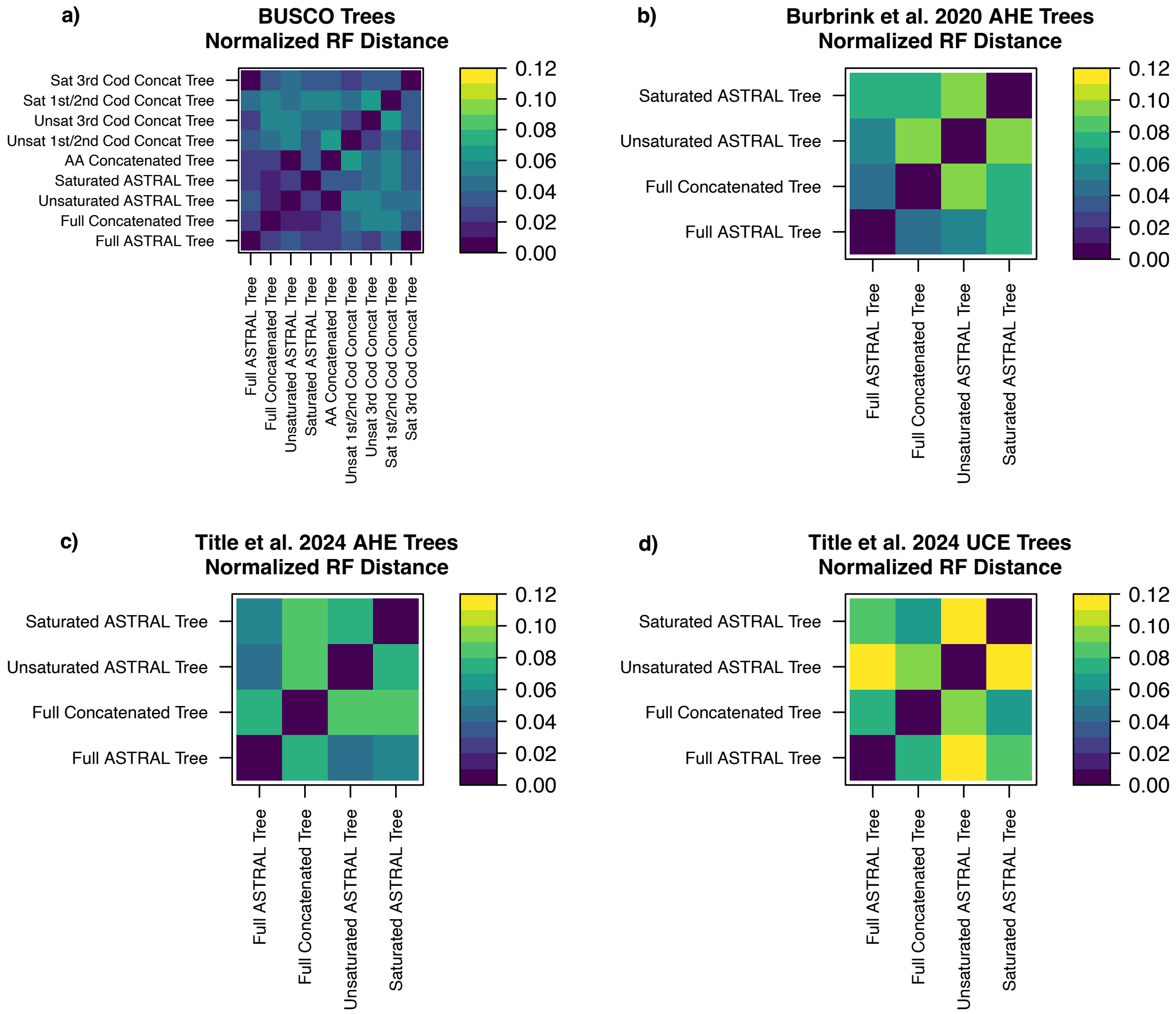

**Supplementary Figure 8: Normalized Robinson-Foulds distance of family-level tree topologies.** Comparison of topological differences between all ASTRAL- and supermatrix-inferred species trees by dataset.

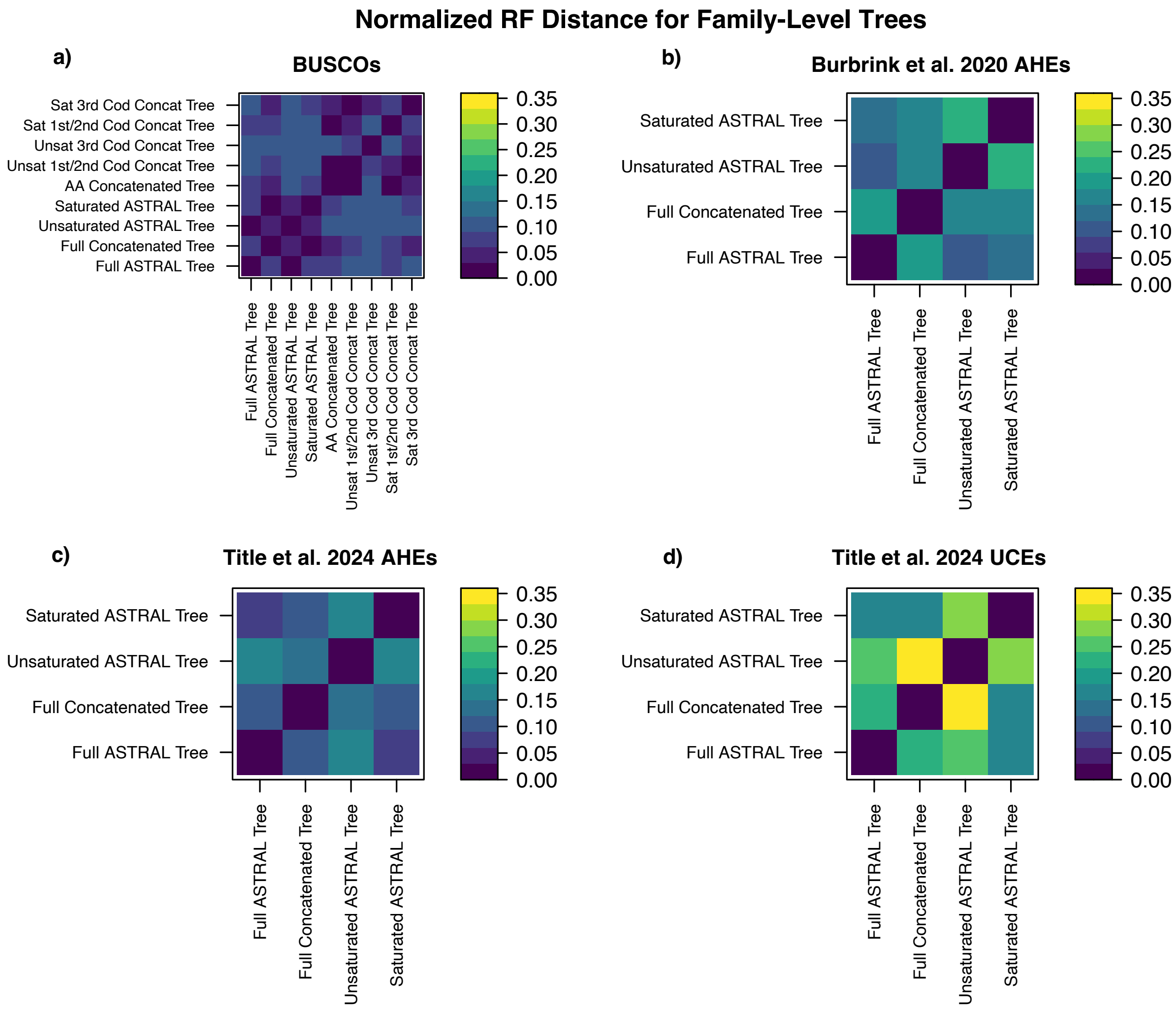

**Supplementary Figure 9: Distribution of concordance factors for full ASTRAL trees.**

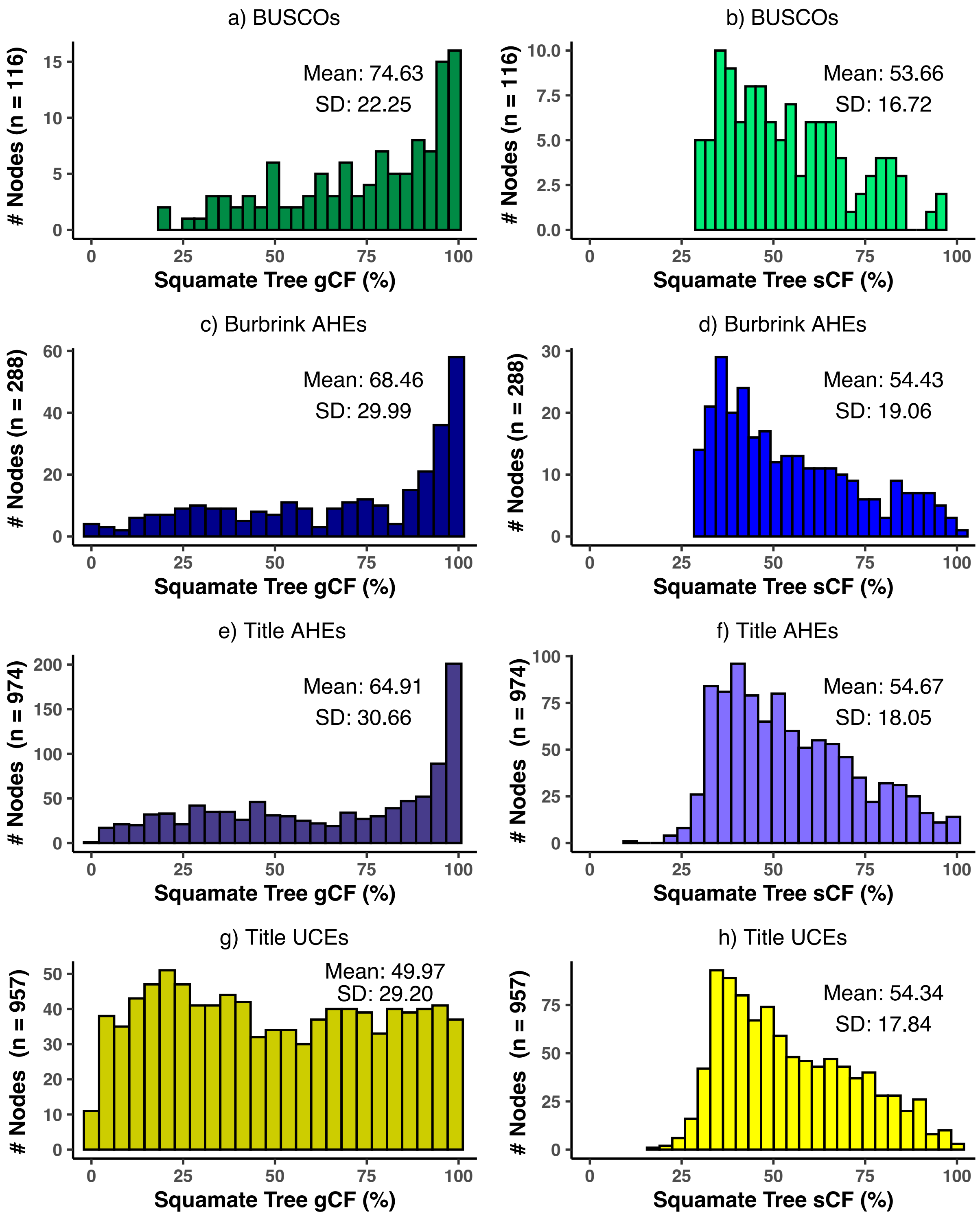

**Supplementary Figure 10: Proportion of saturated alignments by dataset.**

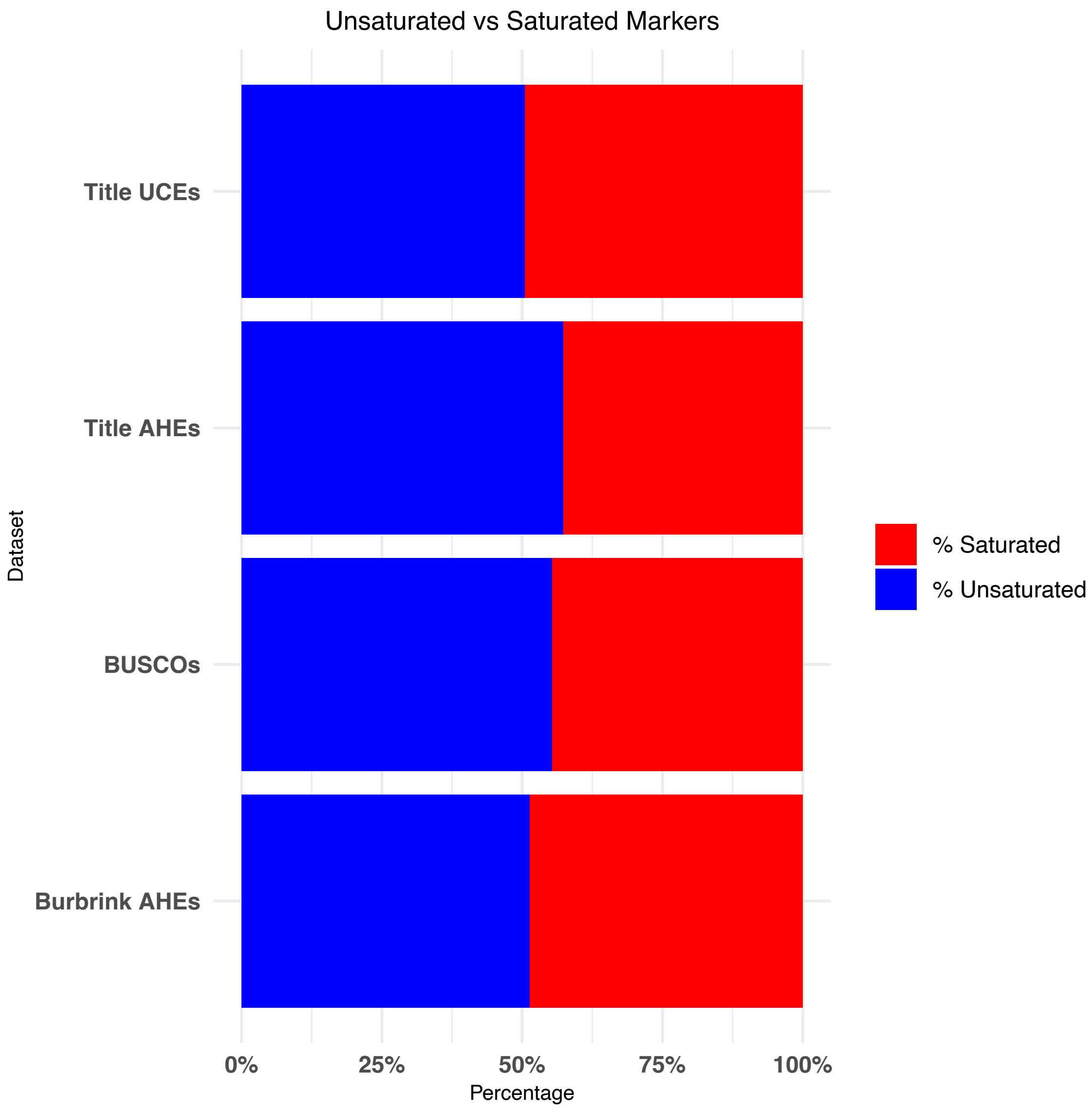

**Supplementary Figure 11: Proportion of saturated alignments by taxonomic family in each dataset**

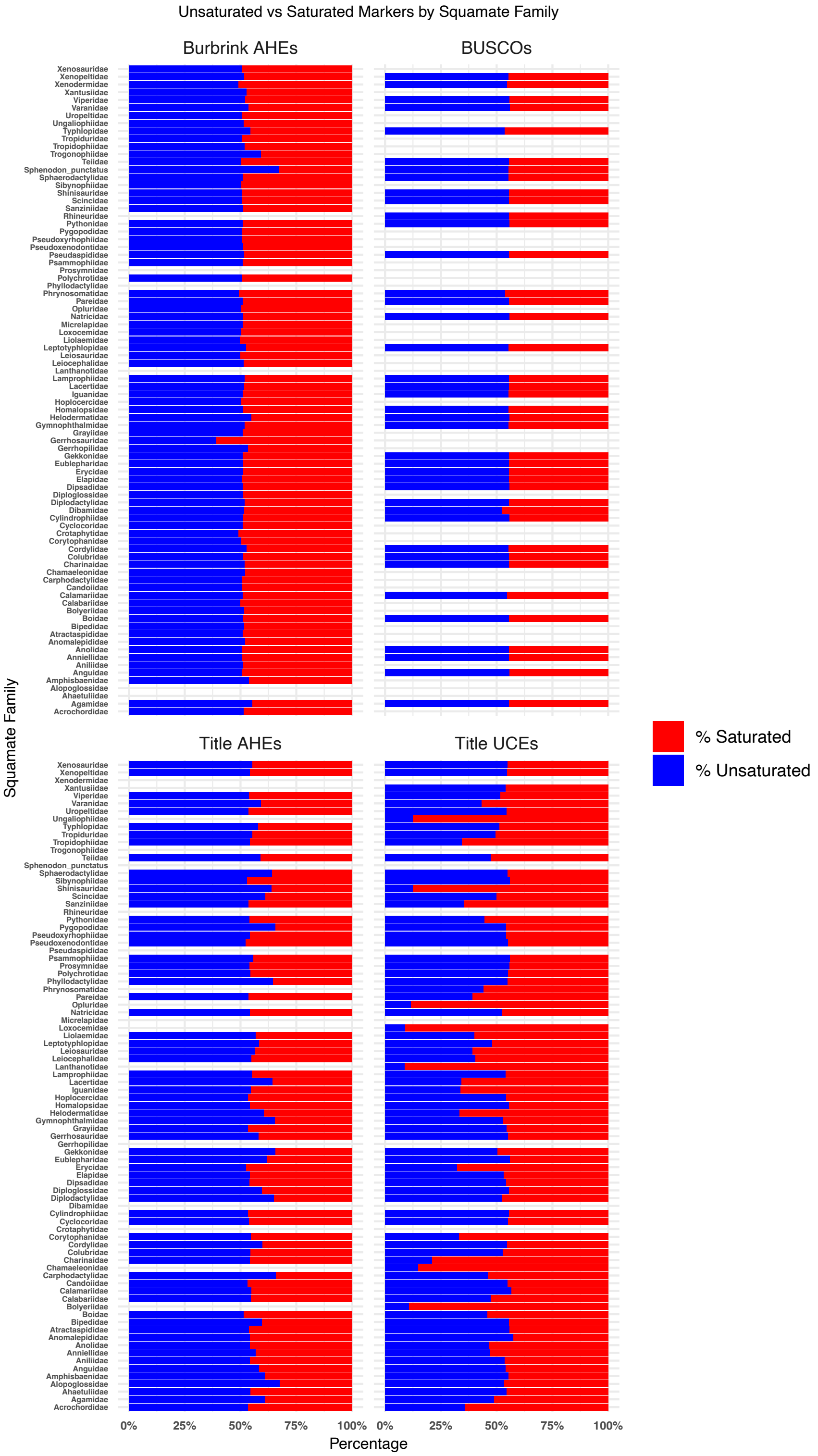

**Supplementary Figure 12: Family-level trees for codon- and saturation-partitioned BUSCO datasets (a-d), and family-level amino acid BUSCO tree (e).**

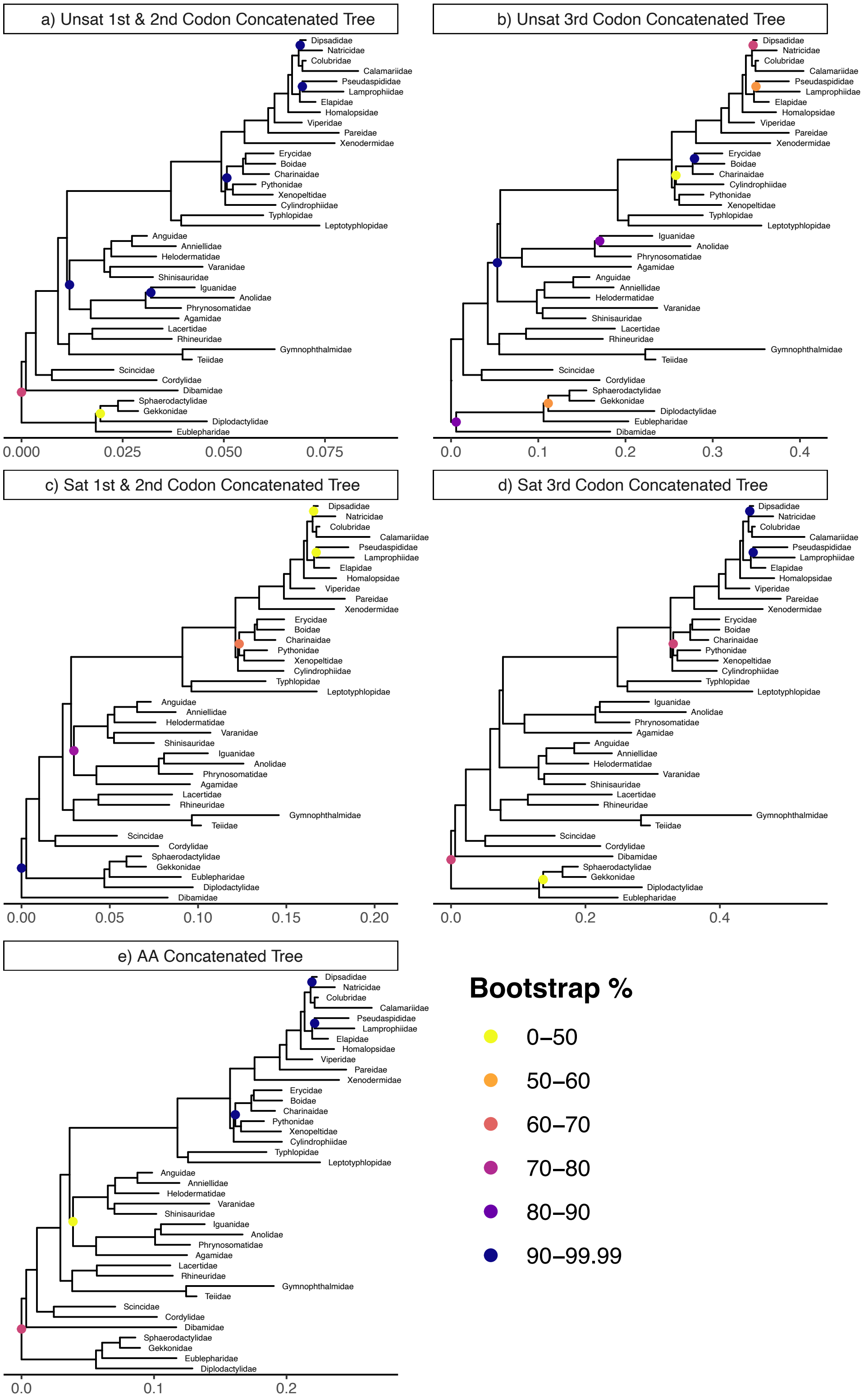

**Supplementary Figure 13: Correspondence analyses results of Synonymous Codon Usage (RSCU).** Based on a concatenated codon-aware alignment of 892 protein-coding genes with 95% taxon sampling by codons (top) and species (bottom).

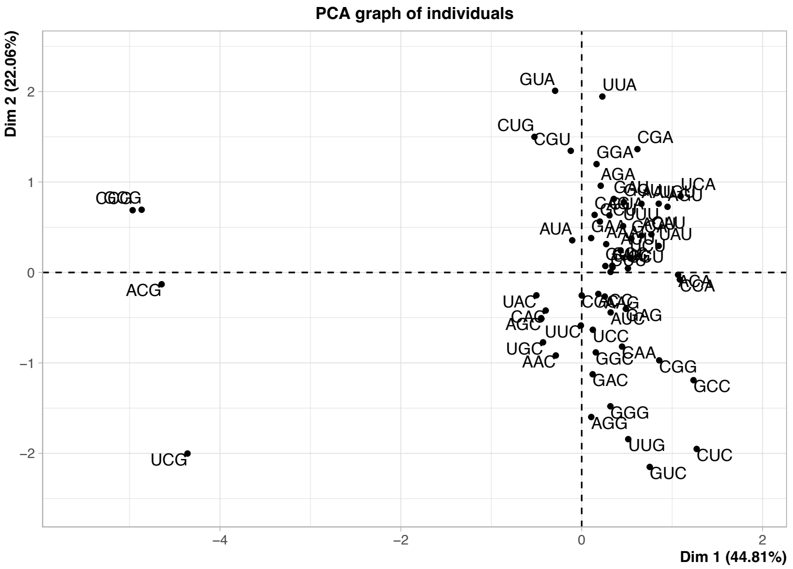


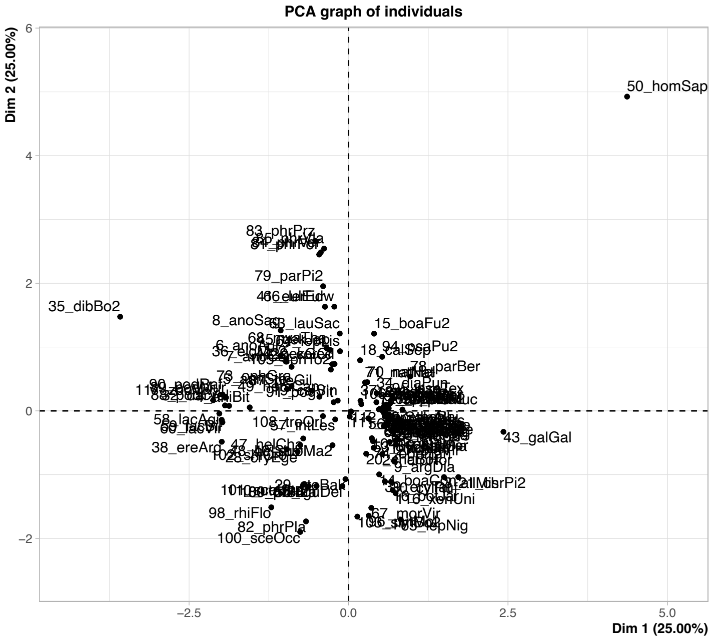

**Supplementary Figure 14: Residual plots from the regressions of observed versus simulated gene concordance factors (gCF).**

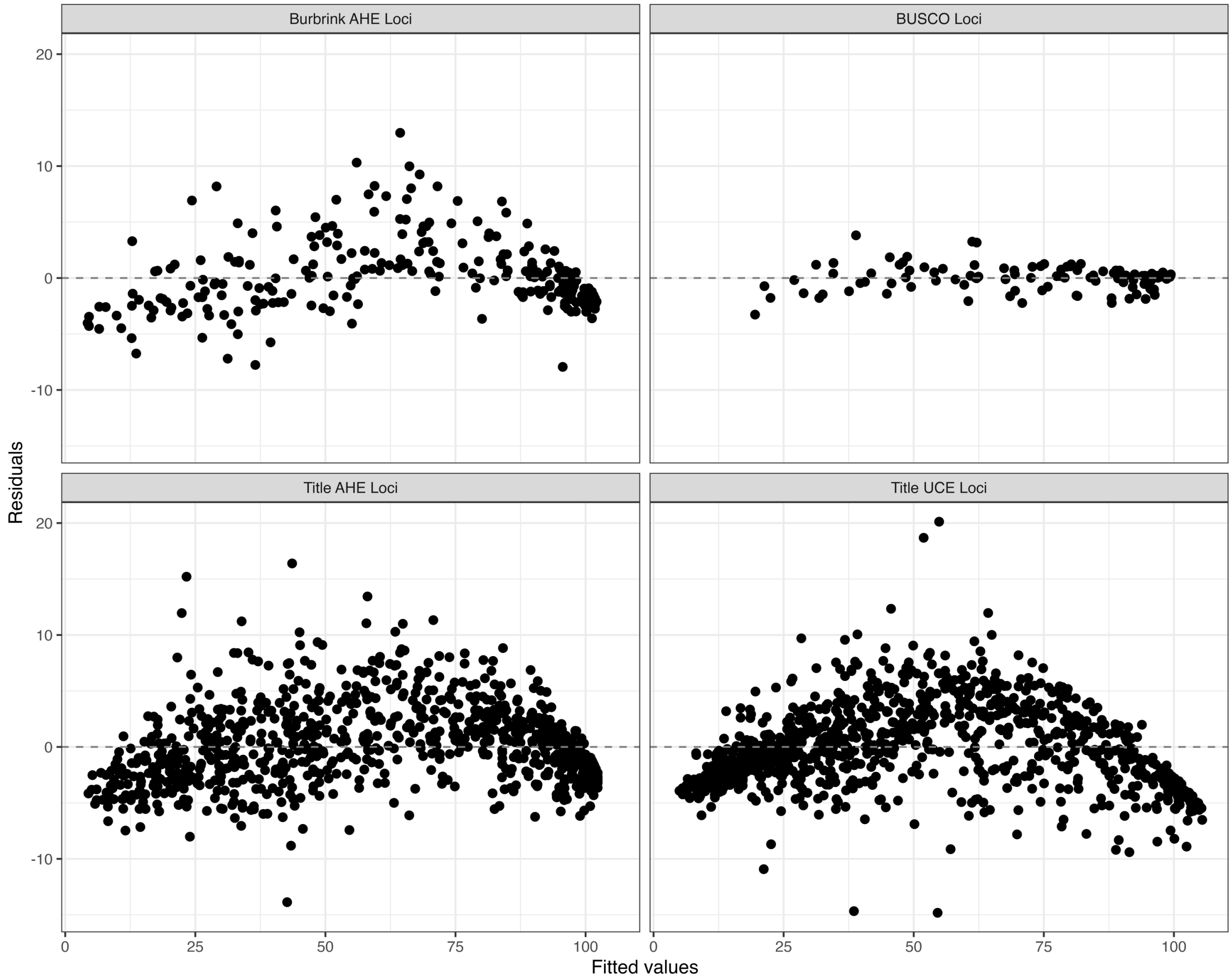

**Supplementary Figure 15. Slopes and intercepts from 1000 iterations of regression tests for observed versus simulated gene concordance factors, by randomly down-sampling each marker set to 100 nodes.** Dashed vertical lines are mean +/- 2 * standard deviation.

### Slopes:

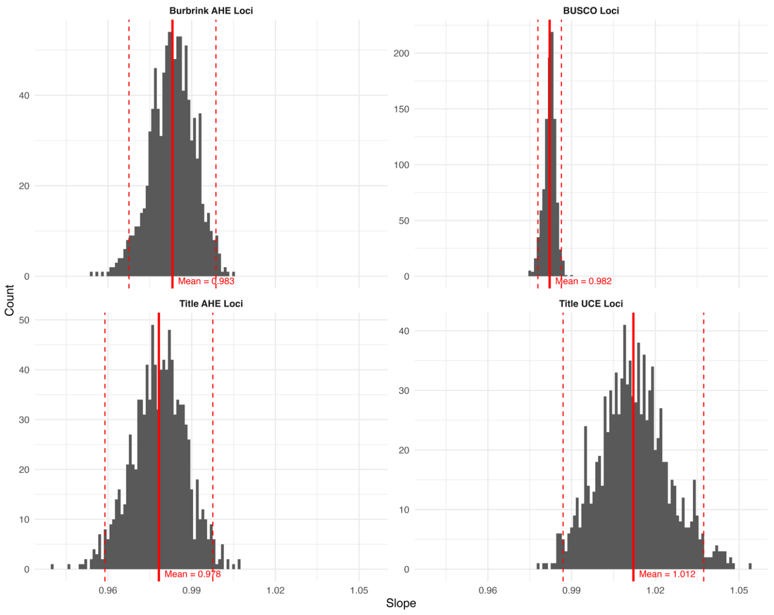


### Intercepts:

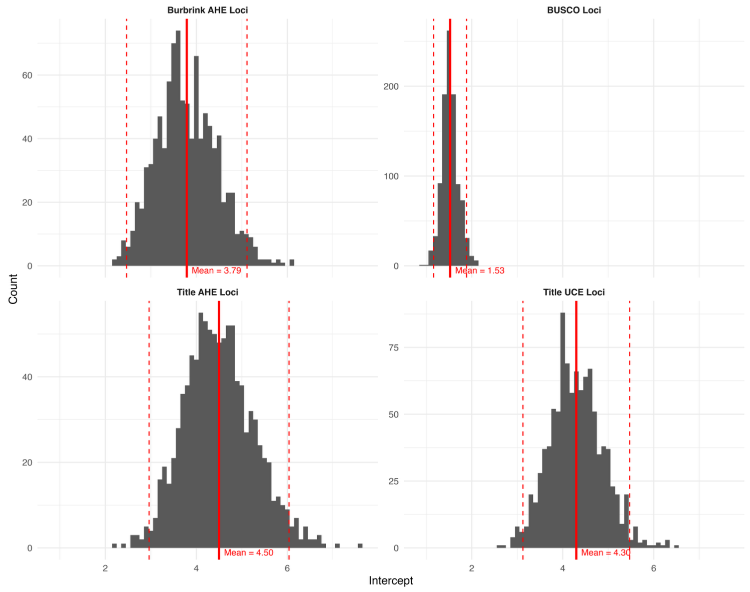
